# Syntaxin 1A Transmembrane Domain Palmitoylation Induces a Fusogenic Conformation

**DOI:** 10.1101/2025.01.24.634780

**Authors:** Dong An, Satyan Sharma, Manfred Lindau

## Abstract

Neurotransmitter release is triggered by the fusion of synaptic vesicles with the plasma membrane, orchestrated by SNARE proteins Synaptobrevin 2 (Syb2), Syntaxin 1A (Stx1A), and SNAP25. Recent experimental studies showed that Stx1A palmitoylation of C271/C272 promotes spontaneous neurotransmitter release. However, the mechanistic role of SNARE transmembrane domain (TMD) palmitoylation in membrane fusion remains unclear.

To investigate the structural and functional implications of TMD palmitoylation, we employed coarse-grained molecular dynamics simulations with the MARTINI force field. In simulations of individual SNAREs and of SNAP-25/Stx1A (t-SNARE) complexes in a membrane the palmitoyl chains of Syb2 and Stx1A localize to the membrane midplane, with Stx1A palmitoyl chains bending toward the extracellular leaflet. Non-palmitoylated Stx1A assumed a conformation where the SNARE domain was lying flat, adhering to the intracellular surface of the membrane. Stx1A dual palmitoylation induced dramatic changes, reducing the tilt of its TMD and stabilizing a more upright conformation of its SND. This conformation resembles the Stx1A conformation in a s Stx1A-SNAP25 t-SNARE complex, providing a potential mechanistic explanation of how Stx1A TMD palmitoylation facilitates early steps in SNARE complex formation and thus promotes spontaneous release.

In simulations of the late steps of layers 5 to 8 SNARE complex zippering in a system of 4 SNARE complexes bridging a 10-nm nanodisc and a planar membrane, FPs spontaneously opened after a few hundred nanoseconds, preceded by distal leaflet lipid transfer and followed by FP flickering conductance before FP closure. At this stage, Stx1A TMD palmitoylation delayed lipid transfer and FP formation and decreased FP flicker open times, whereas the palmitoylation of Syb2 did not affect fusion pore dynamics. These findings suggest that after facilitation of priming before FP opening, Stx1A TMD palmitoylation, directly affects FP dynamics. These results highlight the essential role of SNARE TMD palmitoylation at multiple stages of neurotransmitter release.

**Statement of Significance:** Synaptic vesicle fusion is critical for neurotransmitter release, enabling neuron-to-neuron communication at synapses. Post-translational modifications, such as palmitoylation, are known to influence this process. Using MARTINI coarse-grained molecular dynamics simulations, we examined the impact of SNARE transmembrane domain (TMD) palmitoylation on SNARE protein conformation and fusion dynamics. Stx1A palmitoylation reduces its TMD tilting and changes its SNARE domain conformation, facilitating SNARE complex formation. In fusion pore (FP) simulations, Stx1A palmitoylation delayed FP opening, decreased FP flicker open times, and shortened FP conductance flicker durations by direct interactions with the FP. Interestingly, dual palmitoylation of Stx1A and Syb2 restored flickering duration but decreased FP opening probability within 4 μs, suggesting a nuanced role of TMD palmitoylation in modulating neurotransmitter release.

## Introduction

Neurotransmitter release is a fundamental process in synaptic transmission, involving the fusion of synaptic vesicles with the plasma membrane to transfer chemical signals from the presynaptic to the postsynaptic neuron. This fusion event is tightly regulated and triggered by the arrival of an action potential at the presynaptic terminal (1, 2). The core of the fusion machinery is formed by the SNARE complex, which consists of Synaptobrevin2 (Syb2), Syntaxin1A (Stx1A), and SNAP-25, with additional proteins such as Synaptotagmin1 (Syt1), Complexin, Munc13, and Munc18 playing essential regulatory roles (2, 3).

Post-translational modifications, particularly protein palmitoylation, play a crucial role in modulating the function of key fusion components. Palmitoylation, the covalent attachment of palmitoyl groups to cysteine residues of proteins, has been shown to influence the localization and dynamics of SNARE proteins. For instance, SNAP-25 is palmitoylated at multiple cysteines (C85, C88, C90, C92), enabling its anchoring to the plasma membrane (4, 5). Palmitoylation has also been observed for Syb2 (6). However, a functional role of Syb2 palmitoylation in neurotransmitter release has not been reported.

Recent evidence suggests that in the cell, the Stx1A transmembrane domain (TMD) residues C271 and C272 are both palmitoylated and that this palmitoylation specifically enhances spontaneous neurotransmitter release without significantly affecting evoked release (7). Mutations of these cysteines to valine (C271V/C272V, referred to as CVCV) or inhibition of the palmitoyl acyl transferase in the K260E mutant strongly reduce the frequency of spontaneous release (7), implicating palmitoylation in fine-tuning synaptic activity.

Molecular dynamics (MD) simulations provide a powerful approach for elucidating the molecular mechanisms underlying membrane fusion. The MARTINI coarse-grained force field, in particular, has been widely used to investigate the behavior of SNARE proteins during vesicle docking and fusion (8-12). More recently, all-atom simulations have shed light on the zippering of SNARE complexes that drive the merging of opposing membranes (13, 14).

In this study, we employed coarse grained MD (CGMD) simulations to unravel the roles of SNARE TMD palmitoylation in regulating membrane fusion. While all-atom simulations offer detailed insights, the MARTINI coarse-grained model was chosen to extend the simulation timescales and capture longer-term dynamics while retaining key chemical details (15, 16).

We used two simulation approaches to explore the impact of palmitoylation. In one approach we performed simulations of individual SNAREs or Stx1A-SNAP-25 (t-SNARE) complexes in a planar membrane comparing the properties of non-palmitoylated and palmitoylated Syb2 (palm Syb2) and non-palmitoylated and palmitoylated Stx1A (palm Stx1A). For Stx1A only the dual palmitoylated C271 and C272 was studied because physiologically both cysteines are palmitoylated (7). This approach revealed that the palmitoyl chains of palm Syb2 and palm Stx1A localized to the membrane midplane, with palm Stx1A’s chains bending towards the extracellular leaflet. Palm Stx1A shows reduced TMD tilting and a more upright Stx1A SNARE domain (SND) conformation that facilitates incorporation into the SNARE complex and which may explain the increase of spontaneous release events by Stx1A palmitoylation. In the other approach we performed simulations of systems involving a 10-nm nanodisc and a planar membrane bridged by 4 SNARE complexes (11). In these systems spontaneous fusion pore (FP) formation occurred after a few hundred nanoseconds, followed by transient FP conductance fluctuations. Stx1A palmitoylation delayed FP opening without significantly altering mean open FP conductance.

Interestingly, Stx1A palmitoylation shortened the duration of FP open flicker states and correspondingly decreased FP fluctuations, whereas co-palmitoylation of Syb2 and Stx1A exhibited complex behaviors, with some simulations achieving rapid fusion and others stalling in a hemifusion state. These findings suggest that Stx1A palmitoylation modulates FP dynamics and that cooperative interactions between palm Syb2 and palm Stx1A palmitoyl chains fine-tune these dynamics further. Our results highlight the potential role of SNARE TMD lipidation in modulating the kinetics of neurotransmitter release.

## Material and Methods

### Protein models

The initial condition of the SNARE complex components: Syb2 26-116, Stx1A 189-288 and SNAP25 8-98 (SN1) and 138-206 (SN2), were taken from ref. (11). It is based on the crystal structure 3HD7 with addition of missing residues F287-G288 of Stx1, K83-S98 of SNAP25-SN1 and K201-G206 of SNAP25-SN2 by Modeler (11). The residues C85, C88, C90 and C92 of SNAP25, C271 and C272 of Stx1A and C103 of Syb2 were palmitoylated using the solution builder of CHARMM-GUI (17, 18). The CVCV mutation (C271V and C272V) of Stx1A was also generated via solution builder of CHARMM-GUI (Fig. 1). Subsequently, Syb2 and Stx1A alone and the t-SNARE complex (Stx1A-SNAP25) were converted from the all-atom representation to MARTINI coarse grain models using the modified martinize python script v2.5 by adding the coarse-grained palmitoyl chain representation based on the model of DPPC lipid tails, which also generated the corresponding protein topology .itp files (11, 19). When the Syb2 and the t-SNARE complex (consisting of Stx1A 189-288 and SNAP25 8-98 and 138-206) were converted to MARTINI representations for FP simulations, generating the corresponding protein topology .itp files along with elastic bonds with force constant 1000 kJ/mol/nm^2^ and upper distance bound of 0.9 nm between the backbone beads in the martinize script to facilitate FP formation on microsecond or sub-microsecond timescale. For the t-SNARE complex the same elastic bonds were included between Stx1A, SN1 and SN2 backbone beads to represent a trimer of the complex. Similarly, the t-SNARE complex for t-SNARE complex-plasma membrane simulations, was built as a dimer of Stx1A 189-288 and completed SNAP25, the structure of which was generated from full sequence of SNAP25 (P60881) by ColabFold (20).

**Figure 1.**
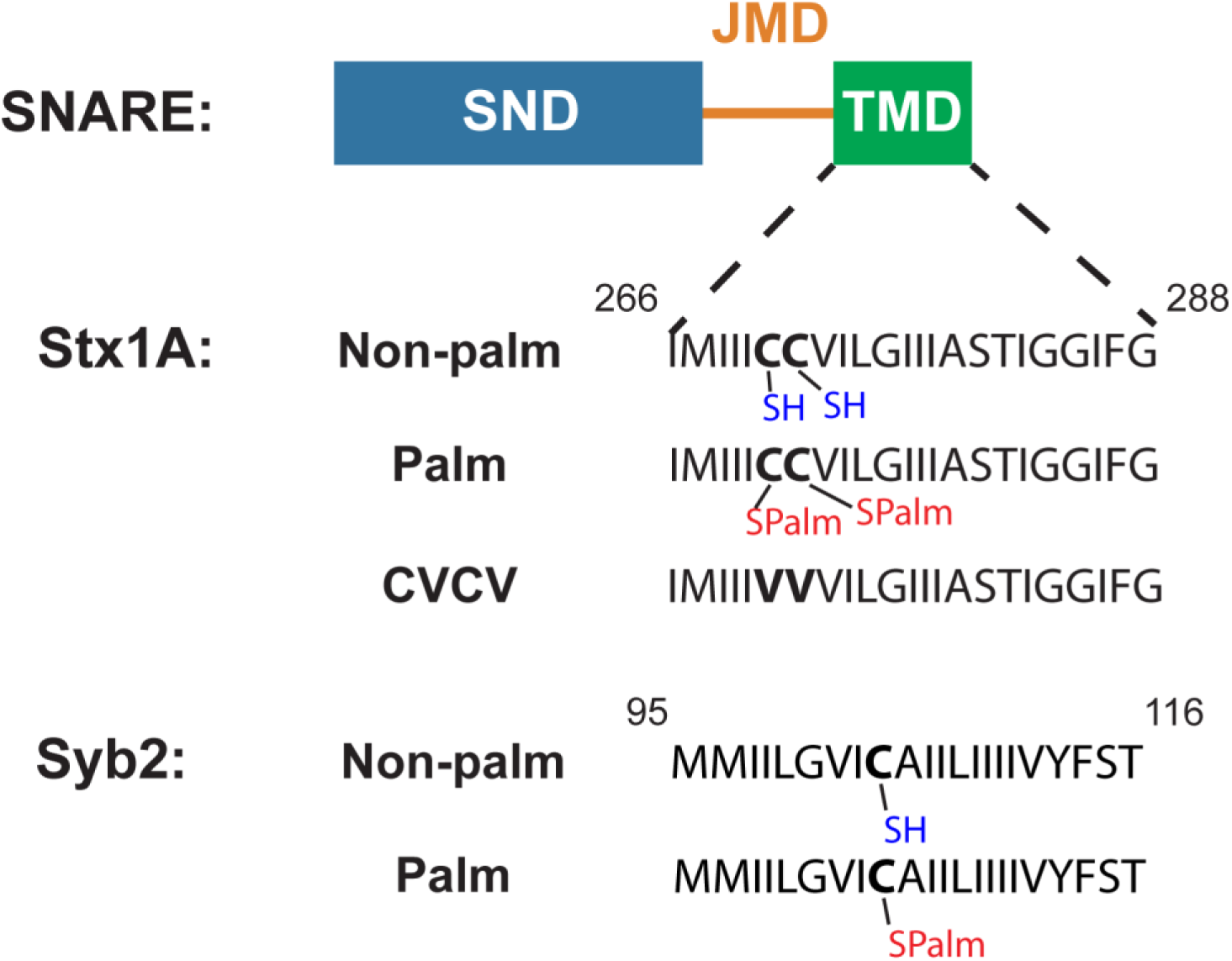
SNARE TMD palmitoylation and mutation in CGMD simulations. Palmitoylation on C271 and C272 for Stx1A and C103 for Syb2 to build TMD palmitoylated SNARE proteins Palm Stx1A and Palm Syb2 were applied to replace the hydrogens at the thiol groups of the side chains of the cysteines to the palmitoyl chain (-COC_15_H_31_). Also, C271V and C272V mutations (CVCV) were also applied to Stx1A to study the Stx1A TMD hydrophobicity in membrane fusion.

### Protein-lipid simulation system preparation

Single copies of Stx1A, palm Stx1A, Stx1A^CVCV^ (Fig. 1) or the corresponding t-SNARE complexes were inserted into an asymmetric plasma membrane via the self-assembly procedure in ref. (21). For non-palmitoylated and palm Syb2, the same procedure was performed but using synaptic vesicle (SV) membrane lipid composition. The intracellular lipids and the extracellular lipids whose components are given in ref. (11) for plasma membrane and in ref. (21) for SV membrane, were placed in two separate 5 nm high boxes with random lipid orientations. Next, the two boxes were combined with 0.5 nm overlap in z dimension. Then the combined box was placed in a 20 nm high simulation box with the protein or protein complex at the center of the box. The x,y positions of the protein or the complex were restrained during membrane self-assembly. After a 500-step energy minimization, the systems were equilibrated in three steps. First a run with 1 ns duration and a timestep of 0.002 ps, restraining the x,y positions of the protein or the protein complex with a force constant of 1000 kJ/mol/nm^2^. The pressure coupling was isotropic since the lipid molecules were oriented randomly in the initial condition. The second step of equilibration with a timestep of 0.02 ps was run for 20 ns, restraining the x,y positions of the protein or the complex with a force constant 100 kJ/mol/nm^2^. In the final step of equilibration, the pressure coupling was changed to semi-isotropic since after the first two equilibration steps, the lipids were no longer oriented randomly but had formed a bilayer. The timestep was kept at 0.02 ps and ran for different durations *t*_mb_as listed in table 1, restraining the x, y positions of the protein or the complex with a force constant 100 kJ/mol/nm^2^. After membrane self-assembly, the lipids that did not insert into the membrane were removed. For more details about the membrane size and membrane lipid composition, please see table S1.

### FP simulations with Nanodiscs system preparation

For FP simulations, the equilibrated initial conditions of an ∼ 32 × 32 nm^2^ plasma membrane and an ∼ 10 nm diameter nanodisc bridged by 4 ternary SNARE-complexes unzipped up to layer 5 were taken from ref. (11). The SNARE complexes, plasma membrane, the nanodisc and water and ions were saved separately in four different pdb files. To perform Stx1A palmitoylation, the two coarse grained Cysteine residues of the Stx1A were replaced by properly aligned palmitoylated Cysteines in all 4 ternary SNARE-complexes and then added back the plasma membrane by *gmx solvate* command. Finally, the atom coordinates of nanodisc, water and ions were added back into the combined pdb file. If this led to removal of a charged lipid, the number of ions and water in the system was adjusted to maintain the zero net charge. Subsequently, the energy of the system was minimized, and 1 ns equilibration was performed with a timestep of 0.002 ps, followed by the actual simulations.

### Simulation details

All simulations were performed using GROMACS version 2021 (22) and the MARTINI force-field 2.1 (15, 16). Berendsen thermostat (310K) and barostat (1 bar) with coupling constant of 1.0 ps were applied. The neighbor list was set to update every 20 steps with verlet-buffer tolerance of 0.005. In protein-lipid simulation systems, two different random self-assembly procedures were performed with different random seeds in placing lipids and self-assembly equilibration to produce two different initial conditions for each kind of protein or protein complex. For each initial condition, 5 simulations with different random seeds were run for 1-2 μs.

For all FP simulations, 10 runs with different random seeds were performed up to 4 μs. After opening, the FP typically closed permanently at some time point and the simulation was discontinued. For more details, please see the .mdp file in https://zenodo.org/records/14219039.

### Analysis

#### Local membrane normal definition for protein and protein complexes

The local membrane normal of the protein or protein complexes was defined as the normal vector of the flat plane fitted to all the coordinates of PO4 beads of the phospholipids whose distance to the juxtamembrane domain (JMD) and transmembrane domains (TMD) of Stx1A or Syb2 was ≤ 5 nm. The direction of the local membrane normal vector is from the extracellular leaflet (EC) to the intracellular leaflet (EC) for plasma membrane and from the intravesicular (IV) leaflet to the cytoplasm leaflet (CP). This local membrane normal was determined for every 1 ns of protein-lipid interaction simulations and was used in determining the time courses of protein tilt angles and palmitoyl chain tilt angles.

#### Protein domains tilting analysis

The orientations of SNARE domains (SNDs), JMDs, and TMDs of Syb2 and Stx1A are the vectors fitted to the coordinates of all the respective backbone beads of the residues whose ranges were given in Table S2 and pointed from N-terminus to C-terminus. The tilt angles θ_SND_, θ_JMD_ and θ_TMD_ are the angles related to the local membrane normal, as illustrated in Fig. 3A_i_.

**Figure 2.**
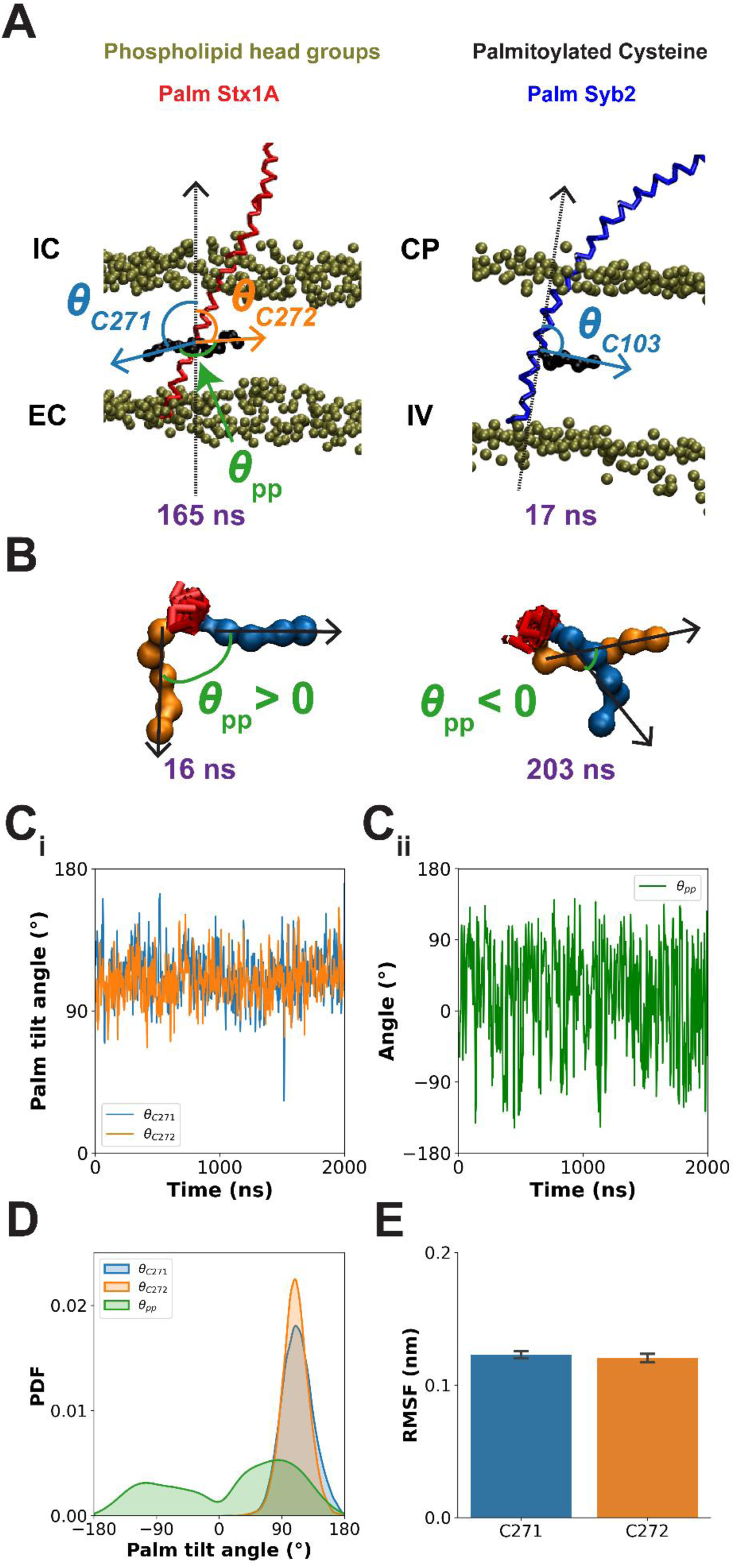
Palmitoyl chain dynamics of Stx1A. (A) Simulation snapshots for the palmitoyl chains of Palmitoyl (Palm) Stx1A and Syb2. The backbones of Palmitoyl Stx1A and Syb2 are colored red and blue respectively. Both the PO4 beads of the plasma membrane (left) and the SV membrane (right) are colored brown and the leaflets for these membranes are labeled. EC: extracellular leaflet, IC: intracellular leaflet, CP: cytoplasm leaflet and IV: intravesicular leaflet. The palmitoyl chains of Stx1A and Syb2 are shown as black chains of beads. The lipid tails are not shown in these snapshots. The simulation times for these snapshots are shown in purple. The vectors of palmitoyl chains for C271 and C272 for Stx1A and C103 for Syb2 are depicted as colored solid arrows and the normal vectors are depicted as the black dash arrows. θ_*C*271_, θ_*C*272_ and θ_*C*103_ ranging from 0° to 180° are the tilt angles of palmitoyl chains for C271, C272 and C103 in Stx1A and Syb2 defined in the method section. This angle > 90° means that the palmitoyl chain bent toward the EC or IV leaflet whereas < 90° means that palmitoyl chain bent toward the IC or CP leaflet. θ_pp_ is the angle between the two palmitoyl chains of C271 and C272 in Stx1A. The snapshot for Stx1A was taken from the 4^th^ simulation of the 2^nd^ initial condition and the snapshot for Syb2 was taken from the 3^rd^ simulation of the 1^st^ initial condition. (B) Simulation snapshots viewed from the N-terminus to C-terminus of the Stx1A (whose backbone is colored red) show that the two palmitoyl chains of Stx1A either pointed away from (left panel) each other or crossed (right panel) each other. The palmitoyl chains for C271 and C272 were colored blue and orange respectively in this panel to distinguish them. (C) The time courses of θ_*C*271_and θ_*C*272_ (C_i_) and the time course of θ_pp_ (C_ii_). They are taken from the 1^st^ simulation of the 1^st^ initial condition. Due to the huge fluctuations, these shown time courses are preprocessed by applying rolling average with a time window of 5 ns. (D) Probability density functions (PDFs) of θ_*C*271_, θ_*C*272_ and θ_pp_ are shown. The two palmitoyl chains of Stx1A tend to point toward the EC leaflet rather than the IC leaflet of the plasma membrane. Their tilt angles for C271 and C272 peak at both 109° with FWHM of 50° and 40°, respectively. The two palmitoyl chains of Stx1A points have two peak values of 84° and –105° with FWHM of 115° and 133° respectively. All 20010 frames from all n = 10 simulations were concatenated to calculate the distributions via kernel-density estimation (27). The peak values and the HWHM of θ_*C*271_, θ_*C*272_and θ_pp_are also listed in Table S3. (E) The average RMSF of the five side chain beads of palmitoyl chains C271 and C272 in Stx1A. They are 0.123 ± 0.003 nm and 0.120 ± 0.003 nm, respectively (mean ± SEM, n = 10). Errorbars in (E): SEM.

**Figure 3.**
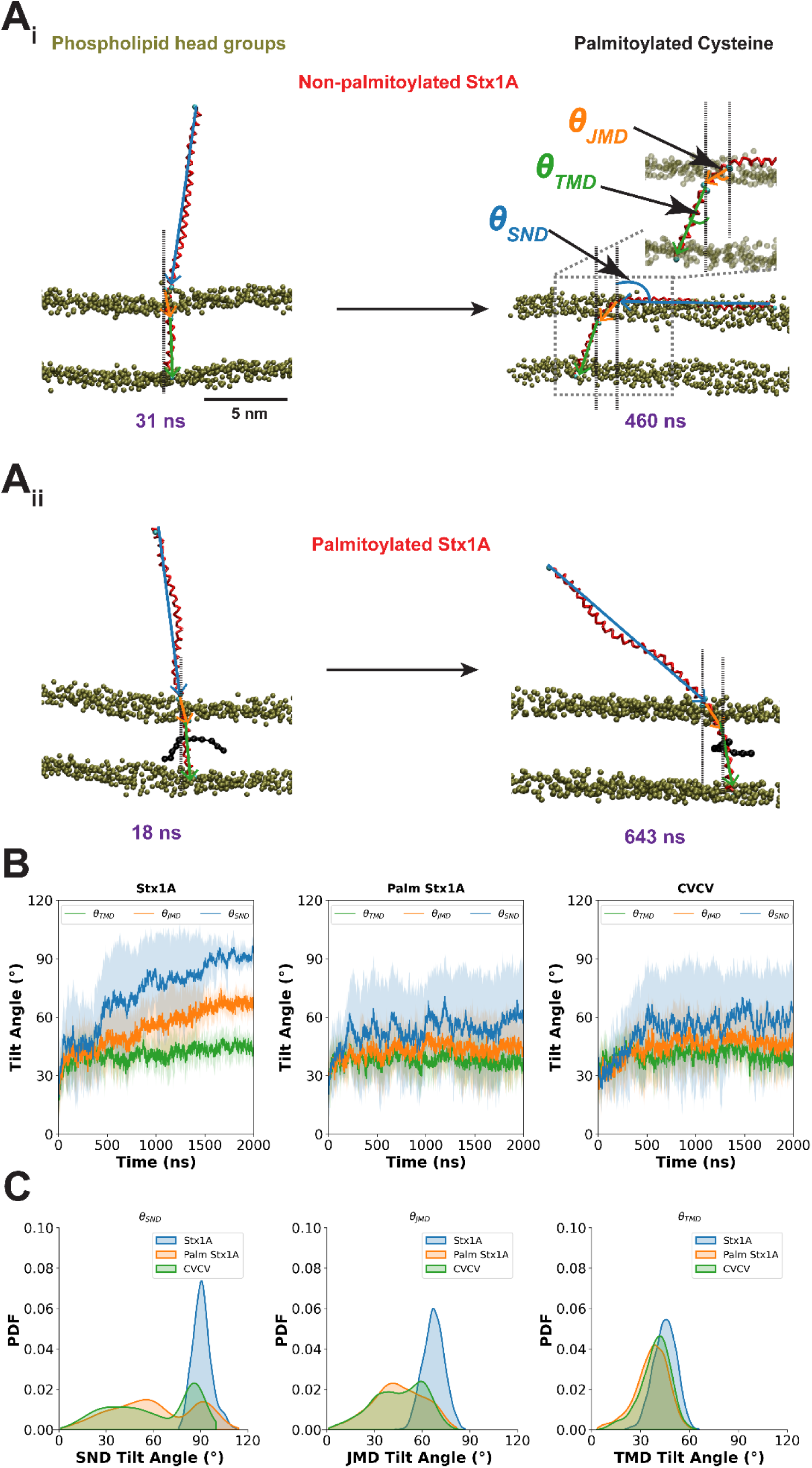
Stx1A TMD palmitoylation and hydrophobicity impairs Stx1A SND, JMD and TMD tilting. (A) Simulation snapshots for non-palmitoylated (A_i_) and palmitoylated (A_ii_) Stx1A (whose backbone is colored red) and a plasma membrane whose phospholipid head groups were colored brown. The zoom in of the region in the dash box was shown but the phospholipid head groups were represented transparent. The lipid tails are not shown in these snapshots. The palmitoyl chains of Stx1A are shown as black chains of beads. The simulation times for these snapshots are shown in purple. The vectors of SND, JMD and TMD for Stx1A are defined as the blue, orange and green arrow fitted to all the coordinates of the backbone beads of these domains. The backbone beads of the residues at the boundaries of these domains are colored cyan. For the information on the initial and the last residues, please see Table S2. The directions are from N-terminus to C-terminus. The angle (range from 0° to 180°) between the vectors and the local bilayer normal (black dash lines) are the tilt angles of these domains (θ_SND_, θ_JMD_ and θ_TMD_), respectively. The snapshots for Stx1A and Palm Stx1A were both taken from their 1^st^ simulation in the 1^st^ initial condition. (B) The average time courses of θ_SND_, θ_JMD_ and θ_TMD_ for Stx1A, Palm Stx1A and CVCV Stx1A are shown (n=10). Shade regions: mean ± SD. The SND of Stx1A ‘lie down’ and irreversibly adhered to the plasma membrane within 2 μs whereas Palm Stx1A typically reached a stable value rather than adhered to the plasma membrane. (C) The probability density function (PDF) of θ_SND_, θ_JMD_ and θ_TMD_ from n = 10 simulations are shown. Frames from the last 200 ns for all simulations were used. The distribution of θ_SND_ of Stx1A peaked at 90° with a FWHM of 12° whereas the θ_SND_ distributions for both Palm Stx1A and CVCV Stx1A are double peaked at 91° and 55° with FWHMs of 22° and 46° for Palm Stx1A, and at ∼ 86° and ∼ 36° with FWHMs of 18° and 83° for CVCV Stx1A. θ_JMD_ distribution of Stx1A is 67° with an FWHM of 16°. CVCV Stx1A θ_JMD_ distribution peaked at 59° and 39° with FWHMs of 42° and 48°, respectively, but Palm Stx1A θ_JMD_distribution only peaked at 41° with an FWHM of 44°. The distribution of θ_TMD_ for Stx1A peaked at 46° with an FWHM was 18° whereas θ_TMD_ distribution for Palm and CVCV Stx1A were peaked at 39° and 42° with FWHMs of 22° and 18°, respectively. For more information, please visit Table S4.

#### Palmitoyl chains tilting analysis

In bilayer simulations, the tilt angles of the palmitoyl chains θ_*C*271_, θ_*C*272_, θ_*C*103_, θ_*C*85_, θ_*C*88_, θ_*C*90_ and θ_*C*92_ are defined as the angle between their vectors connecting the SC1 bead and the center of mass of SC3, SC4 and SC5 beads and the membrane normal as illustrated in Fig. 2A and Fig. 4A. Angles < 90° means that palmitoyl chains tilt toward the IC or the CP leaflet side of the planar membrane whereas angles > 90° means that palmitoyl chains tilt toward the EC or the IV of the planar membrane. For palm Stx1A, the angle between the vectors of the two palmitoyl chains is called θ_pp_. The sign of θ_pp_ is defined in Fig. 2B.

**Figure 4.**
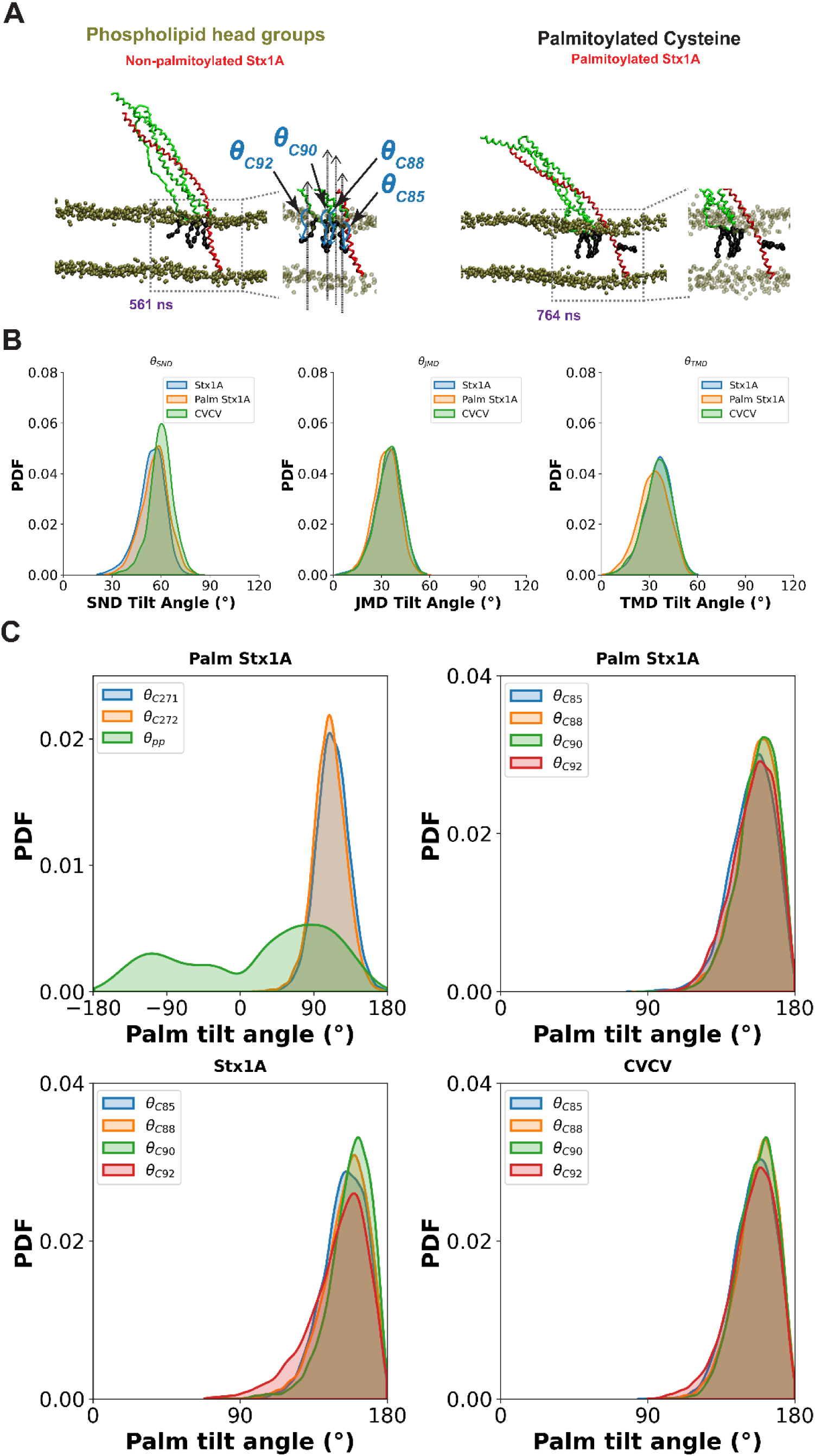
Stx1A in t-SNARE complex did not lie down but maintained a fixed configuration. (A) Simulation snapshots for the t-SNARE with Stx1A and Palm Stx1A in t-SNARE-plasma membrane simulations with palmitoyl chains of C85, C88, C90 and C92 of SNAP-25 and C271 and C272 of Stx1A are shown. Top: Palm Stx1A, bottom: Stx1A. The Stx1A and SNAP-25 are colored red and green respectively. The head groups of the phospholipids in plasma membrane are colored brown. The zoom in of the regions in the dash boxes were shown but the phospholipid head groups were represented transparent. The lipid tails are not shown in these snapshots. Both snapshots of Stx1A and Palm Stx1A were taken from the 1^st^ simulation of the 1^st^ initial condition. The same definition of the palmitoyl chain tilt angles for the chains of C85, C88, C90 and C92 of SNAP-25 (θ_*C*85_, θ_*C*88_, θ_*C*90_, and θ_*C*92_) was applied as in Fig. 2A. (B) The distributions of θ_SND_, θ_JMD_ and θ_TMD_ of Stx1A, Palm Stx1A and CVCV Stx1A in t-SNARE complex defined in Fig. 3A_i_ from n = 10 simulations are shown. The frames of the last 500 ns for all simulations were used. The distribution of θ_SND_ of non-palmitoylated, palmitoylated and CVCV Stx1A peak at 57°, 59° and 60° with a FWHM of 18°, 16° and 14°, respectively. θ_JMD_ distribution of non-palmitoylated, palmitoylated and CVCV Stx1A peak at 37°, 35° and 37° with an FWHM was all 18°, respectively. The distribution of θ_TMD_ for non-palmitoylated, palmitoylated and CVCV Stx1A peak at 36°, 34° and 36° with an FWHM of 20°, 22° and 20°, respectively. For more details, please see Table S6. (C) The distributions of θ_*C*271_, θ_*C*272_, θ_pp_, θ_*C*85_, θ_*C*88_, θ_*C*90_, and θ_*C*92_are shown. All 20010 frames from n = 10 simulations are taken to calculate these distributions. Palmitoyl chains for Stx1A in t-SNARE complex have similar dynamics to those for Stx1A alone. θ_*C*271_ and θ_*C*272_ peaked at 110° and 109° with FWHMs of 45° and 42°, respectively. This is like the values shown in Fig. 2D. θ_pp_peaked at ∼ 87° and –108° with FWHMs of 120° and 137°, respectively. This is also similar to the results shown in Fig. 2D. θ_*C*85_, θ_*C*88_, θ_*C*90_, and θ_*C*92_of SNAP-25 in t-SNARE complex were consistent with all kinds of Stx1A. They peaked at ∼ 154° – 163° with FWHMs of ∼ 28° – 33°. For more information, please visit Table S7.

In FP simulations, the tilt angles θ_*C*271_, θ_*C*272_ and θ_*C*103_ of the palmitoyl chains of palm Stx1A and palm Syb2 are defined as the angles between the palmitoyl chain vectors relative to the vector fitted to the TMD of palm Stx1A and palm Syb2, respectively, rather than the local membrane normal because of the membrane reshaping during FP simulations. The tilt angles for C85, C88, C90 and C92 of SNAP-25 θ_*C*85_, θ_*C*88_, θ_*C*90_ and θ_*C*92_ are relative to the palm Stx1A TMD since SNAP-25 and Stx1A form the t-SNARE complex and SNAP-25 anchors on the plasma membrane. Angles < 90° means that palmitoyl chains tilt toward the N-terminal side of the SNARE TMDs whereas angles > 90° means that palmitoyl chains tilt toward the C-terminal ends of the SNARE TMDs.

#### Palmitoyl chain root mean square fluctuation (RMSF) analysis

The RMSFs for palmitoyl chains are calculated as the average RMSF of each of the five palmitoyl chain beads to quantify the palmitoyl chain structural fluctuation. The palmitoyl chains of C271 and C272 of palm Stx1A in each 1 ns frame were aligned to their structures at 1800 ns in Stx1A-plasma membrane simulations. The palmitoyl chain of C103 was aligned to its structure at 100 ns in palm Syb2-SV membrane simulations. The palmitoyl chains of C271 and C272 of palm Stx1A, and those of C85, C88, C90 and C92 of SNAP-25 were aligned to their structures at 500 ns in t-SNARE-plasma membrane simulations. These alignments removed palmitoyl chain translation and rotation for the RMSF calculation. After alignments, for each bead *i* (*i* = SC1, SC2, SC3 SC4 and SC5), the time-averaged position of the bead 〈***u***_*i*_〉 from the trajectory periods after 1800 ns, 100 ns and 500 ns for Palm Stx1A, Palm Syb2 and t-SNARE planar bilayer simulations was calculated and used as the reference structure. All frames after 1800 ns, 100 ns and 500 ns for palm Stx1A-plasma membrane simulations, palm Syb2-SV membrane simulations and corresponding t-SNARE-plasma membrane simulations were used to calculate the RMSF, respectively. Then, the RMSF of the side chain bead *i* (the square root of the mean square deviation from the time-averaged position of the bead) was calculated as:

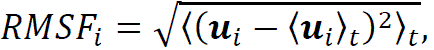

where 〈 〉_*t*_ means time average operation. Finally, the RMSF of a palmitoyl chain is the average RMSF among its five beads.

For the FP simulations, all palmitoyl chains were aligned with the palmitoyl chain structures at 500 ns and all frames after 500 ns were used to calculate the RMSF.

#### Distal leaflet minimal distance analysis

To quantify the intermediate states, the minimal distance between the head groups of any lipids that were initially located at the IV leaflet of the nanodisc and the head groups of any lipids that were initially located at the EC leaflet of the plasma membrane, respectively, was calculated. The cholesterol molecules in this calculation were excluded because of the constant flip flops between leaflets in a membrane. The PO4 bead was chosen as the head group of all phospholipids and GM1 bead was chosen as the head group of all glycosphingolipids. They were used as reference beads for determining the leaflet of the lipids. The midplane of the nanodisc was fitted for the initial frame to the coordinates of all PO4 bead in nanodisc and all individual lipids were assigned to either the IV or the CP leaflet, respectively, according to the PO4 beads being located above or below the midplane. Due to undulations, the plasma membrane was subdivided into 9 (3×3) cells and the assignment of lipids to the IC or EC leaflets was done accordingly for each cell, but taking into account both the PO4 and the GM1 beads (Fig. 5). The minimal distance between any IV and EC head groups was taken to identify the intermediate states. When the minimal distance ≥ 8 nm, the two membranes are separated (Fig. 5A). When the minimal distance is greater than 3 nm but smaller than 8 nm, it is the ‘X’ like stalk state (Fig. 5B). When the minimal distance is ∼ 2-3 nm, the two membranes are hemifused by forming the hemifusion diaphragm (HD) (Fig. 5C, 5D). Finally, when the distance between any lipid head groups from IV and EC leaflet was ≤ 0.8 nm, the first lipid has transferred into the other leaflet (Fig. 5D). For the discontinued simulations, the values from the stopped time to 4 μs were set the same as the last frame of the simulations. The time course of the minimal distance for each simulation was extracted (Fig. 6A). The lag time to the minimal distance smaller than 0.8 nm was extracted as the time of the first leaflet lipid transfer between IV and EC leaflet from each simulation and the corresponding lipid transfer probability *p*_transfer_(*t*) defined as the fraction of simulations that distal leaflet lipid transfer had been occurred within time *t* (Fig. 6C, S5).

**Figure 5.**
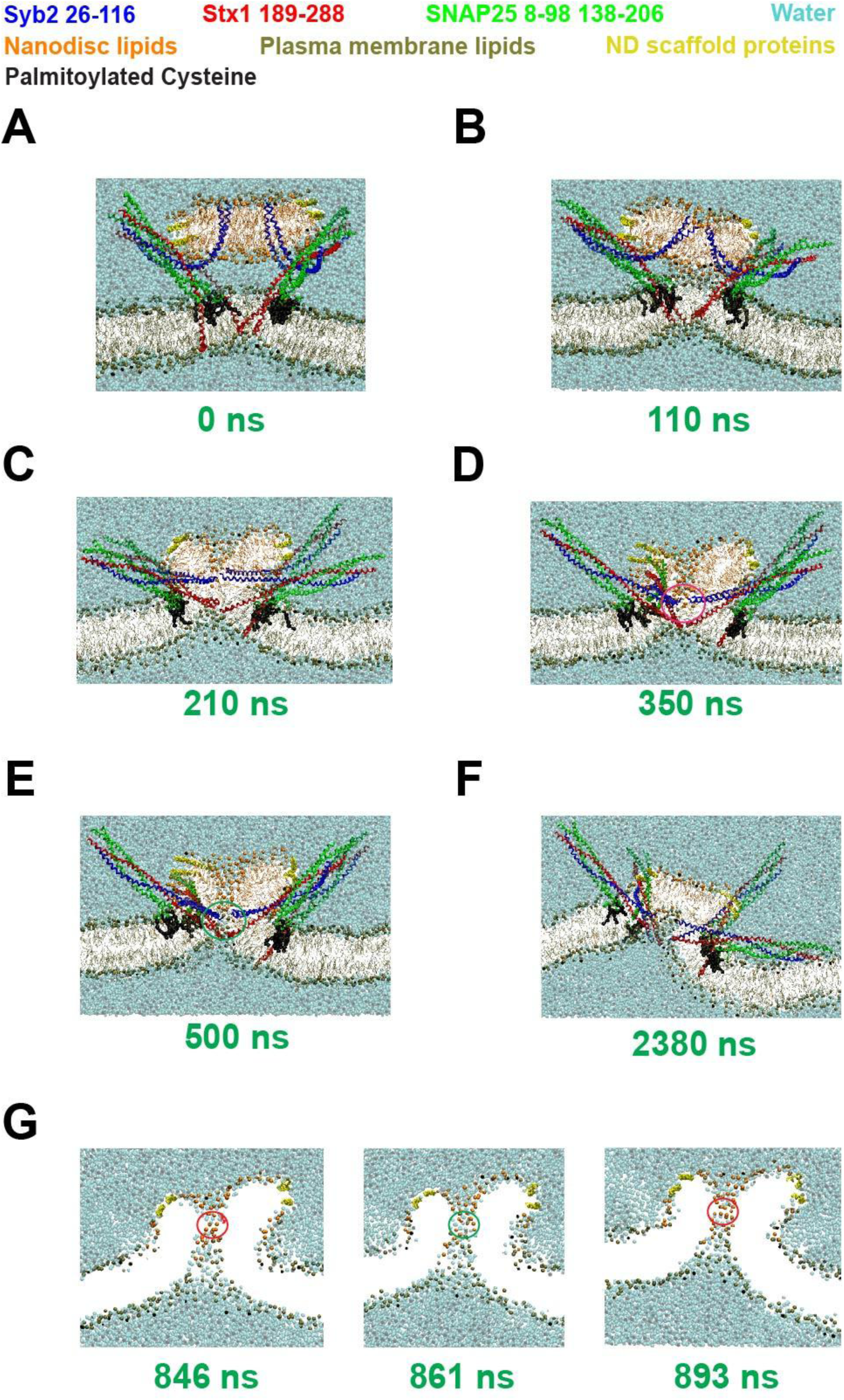
SNARE complexes zippered and governed the membrane reshaping to open FP. Snapshots of a fusion pore simulation with 4 non-palmitoylated SNARE complexes. The numbers in green at the bottoms are the simulation time. The backbone of Syb2, Stx1A and SNAP-25 are colored in blue, red and green, respectively. Water beads are colored in cyan transparently. The lipids of nanodisc and plasma membrane are colored in orange and brown, respectively. The nanodisc scaffold protein was colored in yellow. All snapshots were taken from the 2^nd^ simulation of the non-palmitoylated Syb2 and Stx1A (Syb2/Stx1A). (A) At t = 0, the two membranes stayed away from each other with a distance of ∼ 2 nm. (B) At 110 ns, the CP leaflets contacted with the IC leaflet first, and the head groups of the contacted lipids cleared away from the fusion site by forming an ‘X’ like stalk. (C) At 210 ns, the head groups of the phospholipids from the IV leaflet were closer to that of the phospholipids from the EC leaflet with a minimal distance of ∼ 2 nm. This formed an HD at the fusion site. (D). At 350 ns, the head group of a phospholipid at the EC leaflet transferred to the IV leaflet at the site that contact Stx1A TMD (red circle) across the HD, but the FP had not opened. (E) At 500 ns, the HD disappeared via FP formation and water molecules diffused across the FP marked the green circle. (F) At 2380 ns, This FP closed permanently. (G) FP open flicker during an FP simulation. In this panel, only phospholipid head groups and water beads are shown. At 846 ns, the water molecule was obstructed by phospholipid head groups (red circle) and then the FP fluctuated, and water was allowed to pass through the FP marked by the green circle (an example at 861 ns was shown at the middle). Then the obstruction appeared at 893 ns. This obstruction shown at the red circle typically lasted 1-2 ns.

**Figure 6.**
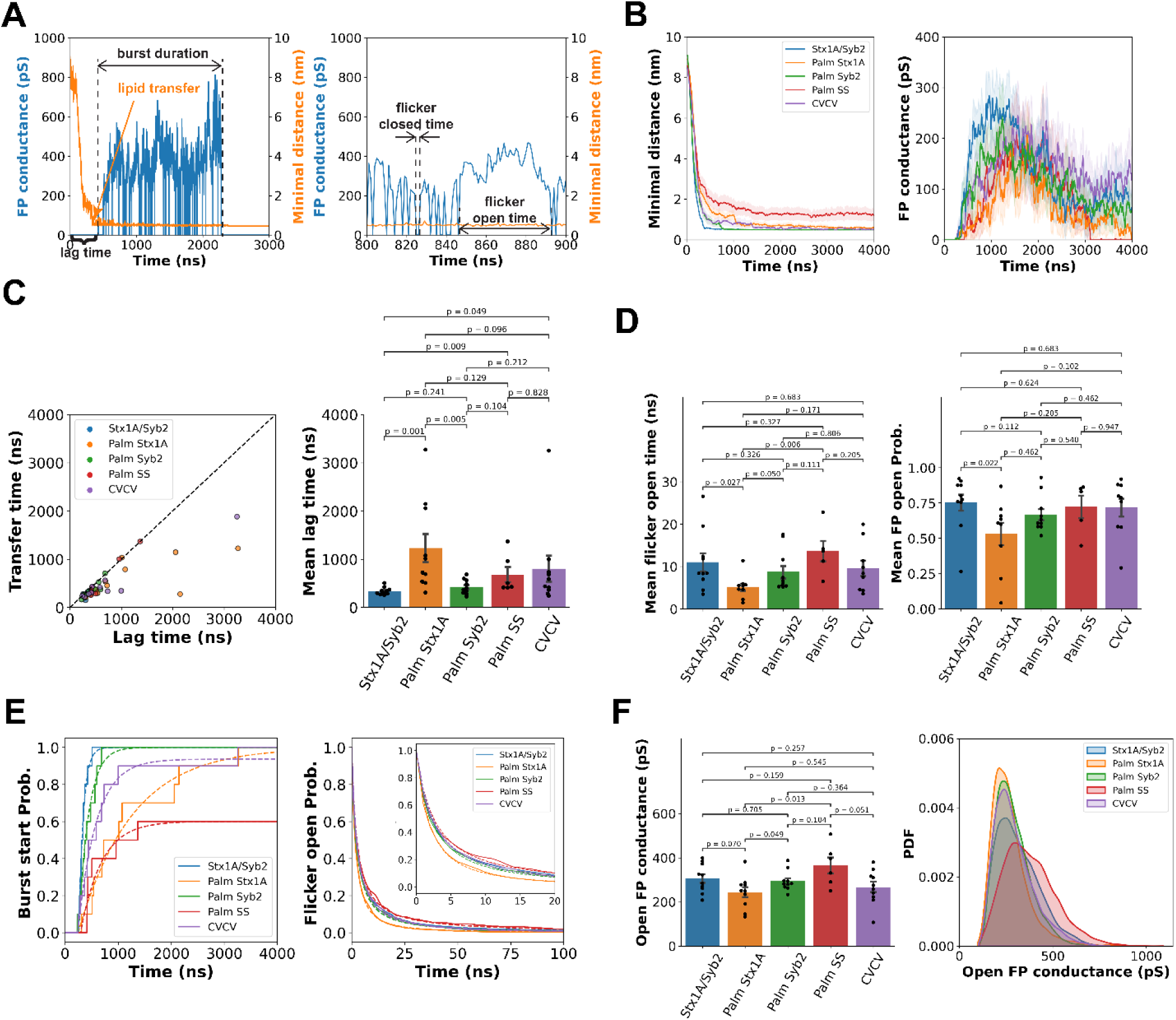
Stx1A palmitoylation delays lipid transfer and FP opening and restricts FP flickering. (A) An example of FP conductance (blue) and distal leaflet minimal lipid head group distance (orange) versus time in a simulation with non-palmitoylated SNARE complex. The burst starts shortly after distal leaflet lipid transfer. The lag time was defined as the delay time from the start of the simulation to the first time when the FP conductance is greater than 1 pS (left panel). The burst duration is the FP lifetime from the lag time to the time when the FP is permanently closed (left panel). The open/closed flicker times are defined as the time intervals when the pore conductance is continuously greater/smaller than 1 pS (right panel). (B) Average time courses of minimal distance between any two phospholipid head groups of EC and IV leaflets (left) and the average time courses of FP conductance (right) showing that the bursts start after a lag time. TMD palmitoylation slowed down the rise of FP conductance by increasing the lag time of FP opening. Shaded area: mean ± SEM. They are smoothed by applying rolling average with a 20 ns time window. (C) The lipid transfer time vs. FP lag time (left) and the mean FP lag time (right). FP lag time is greater than the lipid transfer time. The lag times are Syb2/Stx1A: 340 ± 24 ns, Palm Stx1A: 1234 ± 301 ns, Palm Syb2: 424 ± 48 ns, Palm SS: 680 ± 163 ns, and CVCV: 801 ± 282 ns (mean ± SEM). (D) Mean flicker open time (left) and the mean FP open probability for simulations with Syb2/Stx1A, Palm Stx1A, Palm Syb2, Palm SS, and CVCV SNARE complexes. The mean flicker open times are Syb2/Stx1A: 11 ± 2 ns, Palm Stx1A: 5 ± 1 ns, Palm Syb2: 9 ± 2 ns, Palm SS: 14 ± 3 ns, and CVCV: 10 ± 2 ns (mean ± SEM, n = 10, 9, 10, 5 and 9 by excluding the simulations showed less than 10 flickers). The mean FP open probabilities are Syb2/Stx1A: 0.75 ± 0.06, Palm Stx1A: 0.53 ± 0.09, Palm Syb2: 0.67 ± 0.04, Palm SS: 0.72 ± 0.13, and CVCV: 0.72 ± 0.09 (mean ± SEM, n = 10, 9, 10, 5 and 9 by excluding the simulations showed less than 10 flickers). (E) The cumulative probability distribution of burst start fitted using Eq. 5 (left), and the FP flicker open probability fitted using Eq. 7 (right). The inset is the zoom in for the first 20 ns. The detail parameters are listed in Table S9 and S11, respectively. (F) The mean open FP conductance for with Syb2/Stx1A, Palm Stx1A, Palm Syb2, Palm SS, and CVCV SNARE complexes show no significant difference. However, the distribution of open FP conductance showed that single SNARE palmitoylation and CVCV mutation shorten the range of the pore conductance and inhibit pore fluctuations. But double palmitoylation facilitates fusion pore fluctuation. The black dots in (C, D and F) are individual value of each simulation for each variant. Error bars in (C, D and F): SEM and the first 3 digits of p-values for any pair of comparison using Kruskal-Wallis test are shown. Open FP conductance for all SNARE complexes are Stx1A/Syb2: 306 ± 21 pS, Palm Stx1A: 243 ± 23 pS, Palm Syb2: 294 ± 14 pS, Palm SS: 366 ± 38 pS, and CVCV: 267 ± 25 pS. Note: The 4 simulations for Syb2/Stx1A dual palmitoylation that did not show FP were excluded in these calculations (n = 10 simulations in (C, D, F) except n = 6 for Palm SS). The distribution of Syb2/Stx1A open FP conductance peaked at ∼ 248 pS with the FWHM of ∼ 258 pS (The frame number n = 16283), that of Palm Stx1A are peaked at ∼ 215 pS with the FWHM of ∼ 163 pS (The frame number n = 8860), that of Palm Syb2 are peaked at ∼ 237 pS with the FWHM of ∼ 193 pS (The frame number n = 14410), Palm SS are peaked at ∼ 295 pS with the FWHM of ∼ 312 pS (The frame number n = 8042), and that of CVCV are peaked at ∼ 240 pS with the FWHM of ∼ 194 pS (The frame number n = 17384).

#### SNARE complex zipper dynamics analysis

The zippering dynamics of a SNARE complex was determined by calculating the distances between the backbone beads of the Syb2 and Stx1A layer residues in layers 5-8. (The four pairs of residues are A74-V244, F77-A247, A81-T251, and L84-A254.) These 4 distances for all 4 SNAREs were calculated every 1 ns. For simulations discontinued before 4 μs, we padded the values of the last frame up to 4 μs and then averaged the time course from all 10 simulations since SNDs rarely continue zippering when the FP was permanently closed (Fig. S6A).

#### TMD residue contacts between Syb2 and Stx1A

To examine specific residue contacts between the TMDs of Syb2 and Stx1A, the contact distances between C103 of Syb2 and C271/C272 (or V271/V272) of Stx1A for cysteine contacts, the TMD contact layer distances between Y113 of Syb2 and S281/T282 of Stx1A (23), and the distances between the C terminal residues S115/T116 of Syb2 and G288 of Stx1A for C-terminal contacts, were determined as the minimum distance between any two backbone beads from the respective residues of Syb2 and Stx1A.

Additionally, the contact distances between these residue pairs, including both backbone and side-chain beads, were calculated as the minimum distance between any two beads from the residues in Syb2 and Stx1A. The average backbone contact distances over time for four SNARE complexes are shown in Fig. S6B. The distributions of these backbone contact distances, along with contact distances that include side-chain beads, are illustrated in Fig. S6C and S6D.

#### FP conductance analysis

The algorithm from ref. (11) was used to perform FP analysis. In 1-ns steps, all water beads whose projected distances on the xy-plane were less than 3 nm to the xy-plane projected center of mass of the four backbone beads of T116 Syb2 residue, as they may be close to the FP, were selected. This cylinder with 3 nm radius was then sliced into slices with a thickness of *s* = 0.45 nm. For each slice, the cross-sectional area

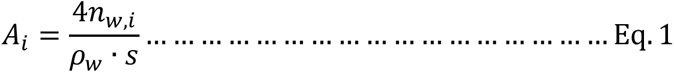

of slice *i* that was filled with water molecules was calculated, where *n*_*w*,*i*_ is the number of water beads in the *i*^*t*ℎ^ slice and ρ_*w*_ = 55.55 mol/L is the water density in real world (4*n*_*w*,*i*_ was used because in MARTINI representation, a water bead represents 4 water molecules (16)). Then we selected all slices with the cross-sectional area *A*_*i*_ calculated from Eq. 1 smaller than 12 nm^2^ (11) since all slices *A*_*i*_ with an area greater than 12 nm^2^ were regarded far away from the FP. Finally, we determined the FP conductance

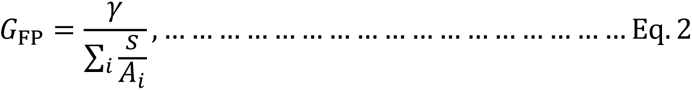

where γ = 1.5 × 10^3^ pS/nm is the bulk conductivity of the solution within the FP, 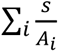 is the sum of the slice width over the cross-sectional area for all the selected slices. For each FP simulation, the time course of FP conductance was extracted (Fig. 6A).

#### FP conductance preprocess and validation

For the discontinued simulations, the FP conductance over the remaining time to 4 μs was set to zero because fusion had occurred, and FPs were closed permanently. For FP simulations which stalled at hemifusion and the snapshots still showed significant conductance with a distal leaflet minimal distance greater than 0.8 nm, the conductance measurements were validated by examining the corresponding snapshots using VMD (24) to identify tilted HD obstructions and the corresponding *G*_FP_(*t*) was set to zero.

#### FP dynamics analysis

The dynamics of all FPs were analyzed by extracting the lag time to burst start, FP flicker open time, and burst duration from *G*_FP_(*t*). The lag time to burst start was defined as the time when the FP conductance was > 1 pS for the first time. The flicker open times are defined as the continuous periods with *G*_FP_ > 1 pS, and the burst duration times are defined as the times from the burst start to the last time with *G*_FP_ > 1 pS within 4 μs. The flicker closed times were defined as the continuous periods with *G*_FP_ < 1 in a burst. The FP open probability is the ratio of the sum of the flicker open times during a burst over burst duration. In the cumulative burst start probability *p*_start_(*t*) curve, the *p*_start_(*t*) is the fraction of simulations that a burst had been initiated at time *t* for each particular SNARE variant. In the flicker open survival curves, all flicker open periods from all simulations of each particular SNARE variant were aligned together to start at *t* = 0, respectively. The flicker open probability *p*_open_(*t*) and the flicker closed probabilily *p*_closed_(*t*) are the fraction of all the aligned flickers remaining open or closed at time *t,* respectively. In the burst duration survival curve, *t* = 0 was set as the time when the burst started, and the burst survival probability *p*_burst_(*t*) is the fraction of the simulations whose FP had not permanently closed at time *t* for each particular SNARE variant.

The open FP conductance was calculated as

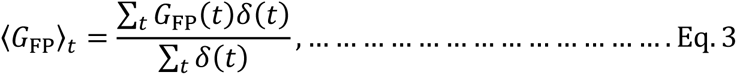

where

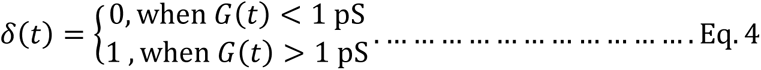

The distributions of *G*_FP_ were calculated from concatenating all frames with *G*_FP_(*t*) > 1 pS from all simulations of a particular SNARE variant together.

#### Statistical tests

Kruskal-Wallis’s test in comparison of any pair of two groups of data was applied since the data distributions were not necessarily symmetrical.

### Model fitting

#### Lag-exponential fitting

The cumulative burst start probability *p*_start_(*t*) and the distal leaflet lipid transfer probability *p*_transfer_(*t*), are fitted using lag-exponential rise function defined as

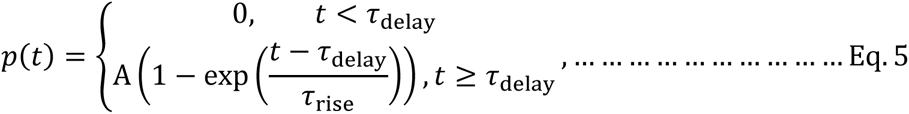

where free parameters τ_delay_ is the delay time that precedes the exponential rise, τ_rise_ is the characteristic rise time of the corresponding probability and *A* is the total change of the probability. The parameters are listed in Table S9 and S10.

The burst survival curve *p*_burst_(*t*) was fitted using lag-exponential decay function defined as

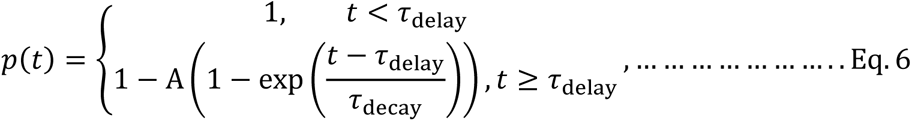

where free parameters τ_delay_ is the delay time preceding the exponential decay, τ_decay_ is the characteristic decay time of the corresponding probability and *A* is the total change of the probability. The parameters are listed in Table S12.

#### Power law fitting

The flicker open cumulative survival probability curves *p*_open_(*t*) and *p*_closed_(*t*) were fitted using the power law defined as

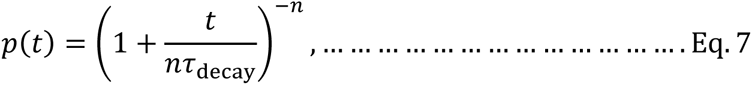

where free parameters τ_decay_ is the characteristic decay time of the corresponding probability and *n* is the exponent factor. The parameters are listed in Table S11.

## Results

### Quantification of Stx1A and Syb2 Palmitoyl Chains in Planar Membrane

To investigate the dynamics of the palmitoyl chains of the cysteines of palm Stx1A and palm Syb2, we conducted molecular dynamics simulations by embedding palm Stx1A and palm Syb2 with palmitoylated TMD into separate 20 x 20 nm^2^ and 15 x 15 nm^2^ planar membrane patches— representing the plasma and synaptic vesicle (SV) membranes, respectively. Notably, the palmitoyl chains attached to C271 and C272 of palm Stx1A, as well as C103 of Syb2, remained predominantly aligned with the midplane of the membranes (Fig. 2A). For more details, please see Movies S1 and S2.

To quantify the behavior of palmitoyl chains of palm Stx1A, the tilt angles (*θ*) of the palmitoyl chains relative to the bilayer normal (Fig. 2A, C_i_) were measured. Correlations between θ_*C*271_and θ_*C*272_ are weak with a Pearson correlation of 0.10. Both, the time courses of a single simulation and the distribution of tilt angles of C271 and C272 palmitoyl chains from all simulations, showed a relatively stable tilt angle of 109° with the full-width at half-maximums (FWHMs) of 50° and 40° for θ_*C*271_ and θ_*C*272_, respectively (Fig. 2C_i_, D), indicating a slight but consistent preferential bend toward the extracellular leaflet of the plasma membrane.

The angle between the two palmitoyl chains (C271 and C272) θ_pp_ is defined as positive when they pointed away from each other or negative when they crossed each other (Fig. 2B, C_ii_). The distribution of θ_pp_ showed two peaks: a peak with a positive value at 84° and the other with a negative angle at –105° with the FWHMs of 115° and 133° respectively (Fig. 2D). Because the area of the peak at 84° is larger than that at –105° (Table S3), the mean angle was 16°, indicating that the two palmitoyl chains preferentially pointed away from each other rather than crossing each other due to steric repulsion.

Root-mean-square fluctuation (RMSF) analysis further demonstrated the structural fluctuations of the two palmitoyl chains (Fig. 2E). The palmitoyl chains of C271 and C272 showed a similar RMSF of 0.123 nm and 0.120 nm, indicating similar structural fluctuations for the two palmitoyl chains when they were buried into the plasma membrane.

The distribution of the palmitoyl tilt angle θ_*C*103_ of Syb2 showed a peaked at 91° with the FWHM of 40°, indicating that the palmitoyl chain of Syb2 stayed mostly perpendicular to the bilayer normal with a similar fluctuation as the palmitoyl chains of Stx1A (Fig. S1). However, the tilt angle distribution showed that the palmitoyl chain occasionally bending toward and inserting either the CP or IV leaflet (Fig. S1A). This is consistent with the two shoulders of the distribution of the palmitoyl tilt angle θ_*C*103_ of palm Syb2 (Fig. S1B). A triple gaussian fit to the palmitoyl tilt angle distribution indicated that θ_*C*103_ ∼ 54° and θ_*C*103_ ∼ 131° were the metastable angles that the chain temporarily assumed. The RMSF of the palmitoyl chain of Syb2 is 0.122 nm, indicating a similar structural fluctuation in the midplane of the SV membrane like the chains of Stx1A. For details of the distribution peak positions and FWHMs, please see Table S3.

### Impact of Stx1A TMD Palmitoylation and Hydrophobicity on Protein Tilt Angles; Syb2 Palmitoylation Shows No Effect

The influence of palmitoylation on the dynamics of SNARE proteins by comparing the tilt angles of the SND, JMD, and TMD relative to the bilayer normal (θ_SND_, θ_JMD_ and θ_TMD_) between Stx1A, palm Stx1A, Syb2 and palm Syb2 were further examined (Fig. 3A_i_, 3A_ii_, S2). To investigate the role of Stx1A TMD hydrophobicity in protein tilting dynamics, the protein domain tilting angles of the C271V and C272V double mutations (CVCV) of Stx1A, which were also used in ref. (7), were also quantified.

In the beginning of the simulations, Stx1A exhibited a sudden increase in tilt angles across all three domains, after which the TMD stabilized in a relatively fixed orientation with the SND lying flat on the plasma membrane surface (Fig. 3A_i_ right, 3B), irreversibly adhering to the membrane surface (θ_SND_ ∼ 90° with an FWHM of 12°). (see Movies S3 for more details). However, the distributions of tilt angles revealed that Stx1A palmitoylation strongly reduced the probability of the SND “lying down” on the membrane surface, and decreased TMD tilt angles (Fig. 3A_ii_, 3B, 3C). Only in 3 out of 10 simulations of palm Stx1A the SND adhered to the membrane within 2 μs. This is consistent with the double-peak distributions of θ_SND_ for palm Stx1A with a strongly reduced peak at 91° and the appearance of a broad peak at 55°with FWHMs of 22° and 46°, respectively. The θ_JMD_ distribution of palm Stx1A only peaked at 41° with FWHM of 44°, and 26° less tilted than Stx1A. The distribution of θ_TMD_for palm Stx1A peaked at 39° with the FWHM of 21°, which is 7° less tilted than Stx1A. For more details, see Table S4.

The CVCV mutant, which was used experimentally as non-palmitoylated Stx1A, also showed a reduced peak at 86° and a broad peak at 36° of θ_SND_ distribution. The distribution of θ_SND_ peaked at 86° and 36° with FWHMs of 18° and 83°, respectively. The change was therefore intermediate between Stx1A and palm Stx1A. In 5 out of 10 simulations of Stx1A^CVCV^ its SND adhered to the membrane within 2 μs. Also, JMD showed intermediate between Stx1A and palm Stx1A with a double-peak θ_JMD_ distribution peaked at 59° and 39° with FWHMs of 42° and 48°, as well as θ_TMD_ distribution peaked at 42° with the FWHM of 18°, which is 4° less tilted than Stx1A.

The θ_SND_, θ_JMD_ and θ_TMD_ between Syb2 and palm Syb2 revealed that the SND typically did not adhere to the SV membrane within 1 μs and Syb2 palmitoylation had no discernible effect on the tilting fluctuations of any of its domains (Fig. S2 and Movie S2, S4). In only 1 out of 10 simulations for Syb2 and palm Syb2 its SND lied down and attached to the membrane. No significant changes in Syb2 tilting dynamics were observed between Syb2 and Palm Syb2. For more information, see Table S5.

### Stx1A in the t-SNARE Complex Resembles the Configuration of Palm Stx1A

To explore how SNAP-25 binding affects the tilting angles of Stx1A and Stx1A palmitoyl chain dynamics, t-SNARE-plasma membrane simulations with Stx1A, Palm Stx1A, and CVCV Stx1A were performed. During the simulations, the Stx1A tilt angles of all three domains in a Stx1A-SNAP25 t-SNARE complex θ_SND_, θ_JMD_ and θ_TMD_ equilibrated within ∼500 ns (Fig. 4A, S3A; for details, see Movies S5, S6).

In t-SNARE-planar membrane simulations the SND did not tilt down to the membrane surface and Stx1A palmitoylation did not affect the distribution of θ_SND_, θ_JMD_ and θ_TMD_ compared with the complex with Stx1A (Fig. 4B). The distribution of θ_SND_ for palm Stx1A and Stx1A in t-SNARE complex peaked at 57° and 59°, respectively, consistent with the second peak value of θ_SND_in Palm Stx1A-plasma membrane simulations at 55° (Figure 3C, 4B). The distribution of θ_JMD_ and θ_TMD_ for palm Stx1A and Stx1A in t-SNARE complex peaked at ∼ 36°, similar to the peak of θ_JMD_ and θ_TMD_ of Palm Stx1A at ∼ 40°. These results strongly suggested that palmitoylation of Stx1A promotes an orientation that facilitates Stx1A assembly with SNAP-25 in a t-SNARE complex. CVCV mutants did not alter the SND, JMD or TMD tilting dynamics in t-SNARE complex. For details, please see Table S6.

The Stx1A palmitoyl chains in the t-SNARE complex remained similar to Stx1A-plasma membrane simulations, locating in the bilayer midplane, slightly bending toward the EC leaflet (Fig. 2C, 4C, S3B). In contrast, the palmitoyl chains of C85, C88, C90, and C92 in SNAP-25 inserted into the intracellular leaflet of the plasma membrane, stayed parallel to the planar membrane normal and pointed toward the extracellular leaflet as expected (Fig. 4A, C), consistent with the angle distribution peaks at ∼ 160° with FWHMs of ∼ 30°. Neither Stx1A palmitoylation nor the CVCV mutation had a significant effect on the tilting dynamics of these four SNAP-25 palmitoyl chains. For details, please Table S7.

Further RMSF analysis revealed that the palmitoyl chains of Stx1A in t-SNARE complex exhibited greater structural fluctuations (∼ 0.12 nm) compared to those of SNAP-25 (∼ 0.11 nm), similar to the fluctuations without SNAP-25. Neither Stx1A palmitoylation nor the CVCV mutation altered the tilting fluctuations of the SNAP-25 palmitoyl chains (Fig. S3B). This suggested no interaction between palmitoyl chains of Stx1A and SNAP-25. For more information, see Table S8.

### Neither Palmitoylation of SNARE TMDs nor Stx1A TMD Hydrophobicity Alter the Membrane Fusion Pathway

In FP simulations between a 10-nm nanodisc and a ∼ 32 × 32 nm^2^ plasma membrane bridged by four SNARE complexes initially unzippered up to layer 5 (Fig. 5A, S4A), the nanodisc was ∼ 2 nm away from the plasma membrane. Within 110 ns, the CP leaflet of the nanodisc and the IC leaflet made contact, forming an “X”-shaped intermediate stalk (Fig. 5B, S4A) via lipids from IC and CP leaflets stay perpendicular to both the bilayers at the fusion site, whereas the head groups of phospholipids from the IV leaflet had the minimum distance of ∼ 8 nm to those of phospholipids from the EC leaflet. During this process, hydrophilic residues at the C-terminus of Syb2 and Stx1A governed the reshaping of the nanodisc and plasma membranes, respectively (Fig. 5A-F, S4A). Then, in the next 100 ns, the continued zippering of the four SNARE complexes further brought the IV leaflet of the nanodisc close to the EC leaflet, whereas the lipids in the fused CP and IC leaflets started expansion at the fusion site (Fig. 5C, S4A). The hemifusion diaphragm (HD) was formed when the phospholipids from the IV leaflet had the minimum distance of ∼ 2 nm to the phospholipids from the EC leaflet. About 100 ns later, the phospholipids at the EC leaflet transfer to the IV leaflet in the vicinity of the Stx1A TMD without FP opening (Fig. 5D).

Then, after 150 ns, an FP spontaneously opened at the site of HD with more phospholipids at the EC leaflet transferring to the IV leaflet, allowing water molecules to diffuse intermittently across the FP (Fig. 5E) until the FP typically closed permanently within 4 μs (Fig. 5F). The water diffusion underwent a flickering behavior, showing intermittently opening and closing FP flickers, since FP was occasionally obstructed by phospholipid head groups (Fig. 5G, Movie S7).

Simulations incorporating Palm Stx1A, CVCV Stx1A, Palm Syb2, and Syb2/Stx1A dual palmitoylation (Palm SS) followed the same FP opening pathway but changed the kinetics of FP opening and flickering. In all cases, the FP opened in the HD, exhibited flickering water transportation, and ultimately closed permanently (Fig. S4B-D, Movie S8-S11).

### Phospholipid transfer between EC and IV leaflets precedes FP opening

The minimal distances between any pair of lipids from IV and EC leaflets except cholesterol, and FP conductance were quantified to understand the mechanism of membrane fusion and how neurotransmitter release is modulated by FP flickering (Fig. 6A). The minimal distance between any pair of lipids from IV and EC leaflets ≤ 0.8 nm indicated that lipid transfer between the two distal leaflets had occurred. Simulations with palm SS showed no lipid transfer in 4 out of 10 simulations with the minimal distances stalled at ∼ 2 nm, indicating that the lipids remained localized in the respective leaflets of the HD whereas with all other SNARE systems, lipid transfer was observed and an FP opened, and membranes fused within 4 μs. (Fig. 6B). This severe impairment of fusion with palm SS is consistent with a larger stable minimal distance of ∼ 1.2 nm for Palm SS than for the other SNARE variants (∼ 0.5 nm) (Fig. 6C, S5), due to steric interactions between the palmitoyl chains of palm Syb2 and palm Stx1A.

After the start of the simulations, FPs opened spontaneously after a lag period of a few hundred nanoseconds and exhibited flickering conductance before eventually closing again (Fig. 5, 6A, S4 and Movie S7-S11). This flickering phase, characterized by intermittent FP opening and closing, is termed the “burst duration” (Fig. 6A). The average conductance time courses showed that the conductance rise became slower and decreased in amplitude by SNARE TMD palmitoylation (Fig. 6B). For Palm SS, all FPs are closed at ∼ 3 μs. The initial distal lipid transfer time preceded the FP lag time and therefore FP formation (Fig. 6C). The mean delay time between initial lipid transfer and FP formation is 65 ± 21 ns (n = 10) for Stx1A/Syb2. The mean delay times for Palm Syb2 and palm SS are similar to Stx1A/Syb2 (66 ± 22 ns, n = 10 and 73 ± 41 ns, n = 6). For Palm Stx1A and CVCV the mean delay times between lipid transfer and FP formation are much longer, 599 ± 240 ns and 300 ± 135 ns, respectively (n = 10).

### Stx1A palmitoylation delays FP opening and Restricts FP Fluctuations

The palmitoylation of Stx1A significantly increased the FP lag time to 1234 ± 301 ns (mean ± SEM, n = 10) compared to the non-palmitoylated SNARE complex (Stx1A/Syb2) (340 ± 24 ns, n = 10, p = 0.001, Kruskal-Wallis test) (Fig. 6C). The mean lipid transfer time also increased to 634 ± 121 ns (n = 10) from 276 ± 20 ns (n = 10, p = 0.016) (Fig. S5A) and therefore partially accounts for the increase in FP formation lag time. The cumulative burst start probability curves were fitted with a delayed exponential function in Eq. 5 (Fig. 6E, left). Stx1A palmitoylation increased the rise time τ_rise_ by a factor of 12 from 85.8 ± 0.3 ns of Stx1A/Syb2 to 1,011 ± 9 ns but the delay times τ_delay_ were similar (257.8 ± 0.2 ns of Stx1A/Syb2 and 245.3 ± 3.7 ns of Palm Stx1A) (Fig. 6E). Correspondingly, Stx1A palmitoylation increased the rise time τ_rise_of lipid transfer by a factor 8 from 61.0 ± 0.7 ns to 478 ± 3 ns (Fig. S5B) but slightly shortened the delay time from 231.8 ± 0.5 ns to 187.1 ± 1.8 ns.

Stx1A palmitoylation also decreased the mean flicker open time from 11 ± 2 ns of Stx1A/Syb2 to 5 ± 1 ns (n = 10 and 9, p = 0.027) and the mean FP open probability from 0.75 ± 0.06 (n = 10) to 0.53 ± 0.09 (n = 9, p = 0.022). However, the other SNARE variants did not show significant changes in these parameters (Fig. 6D). The survival curves of FP flicker open times (Fig. 6E, right) were not single-exponential but could be well fitted with a power law in Eq. 7

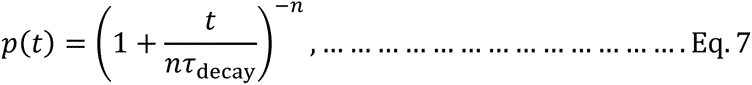

indicating heterogeneity of time constants. For large n the time course becomes exponential with time constant τ_decay_ but with decreasing n the time course becomes increasingly non-exponential with a slow tail. Stx1A palmitoylation shortened τ_decay_ of the flicker open time from 2.4 ns to 1.9 ns and increased the exponent *n* from 1.2 to 1.6.

The flicker closed times are mainly 1 ns across all SNARE variants, mainly due to the occasionally obstruction of FP by lipid head groups. This effect could result from the MARTINI coarse-graining. However, these changes of FP dynamics were partly restored by Syb2 palmitoylation (Fig. 6D, 6E). Despite differences in flickering behavior, the burst durations were similar across all groups, including Stx1A/Syb2, Palm Stx1A, Palm Syb2, Palm SS and CVCV variants. For details, see Tables S9-S12.

Finally, open FP conductance analysis revealed that SNARE palmitoylation did not significantly alter the mean open FP conductance estimated by Eq. 3 and Eq. 4 (Fig. 6F). However, palmitoylation of either Stx1A or Syb2, as well as the CVCV mutation, decreased the peak location of the open FP conductance distributions from 248 pS to 215 pS, 237 pS and 240 pS, and decreased the FWHM from 258 pS to 163 pS, 193 pS and 194 pS, respectively. In contrast, Palm SS restored the peak location to 295 pS and facilitate a greater FWHM of 312 pS (Fig. 6F).

### SNARE zippering dynamics are not affected by SNARE palmitoylation

The dynamics of SNARE complex zippering by measuring the distances between the backbone beads of layer 5-8 residues of Syb2 and Stx1A showed that all SNARE complexes underwent rapid zippering, with the distance between the backbone beads of these residues stabilizing within < 1 μs (Fig. S6A). The average zippering time courses for all SNARE systems are similar. One small difference is that when Stx1As are palmitoylated, the distances between the layer residues at the layer 7 and 8 are ∼ 0.70 nm compared with ∼ 0.80 nm for non-palmitoylated Stx1A systems. Interestingly, even if in the 4 out of 10 simulations for Stx1A/Syb2 dual palmitoylation stalled at hemifusion, their SNAREs were fully zippered but without leading to FP formation. The Stx1A^CVCV^ did not affect SNARE zippering dynamics.

### Stx1A and Syb2 TMD palmitoyl chains contact each other and reduce the interactions between Stx1A and Syb2 TMDs

Analysis of the contact distances between the backbone beads of SNARE TMD cysteines (Syb2 C103-Stx1A C271/C272), TMD layer residues (Syb2 T113-Stx1A S281/T282) and C-terminal residues (Syb2 S115/T116-Stx1A G288) revealed that SNARE TMD palmitoylation slowed down the formation of contacts between these residue pairs compared to Stx1A/Syb2 (Fig. S6B).

The distributions of the backbone bead contact distances of the SNARE TMD cysteines showed that the palm SS shifted the highest peak from 1.0 nm to 1.2 nm (Fig. S6C). In addition, the peaks at 0.76 nm for Stx1A/Syb2 and 0.85 nm for Palm Syb2 disappeared for Palm Stx1A and Palm SS. Palm SS even showed the second highest peak at 1.7 nm.

The contact distances including the cysteine side chains (side chain contacts) showed a main peak at 0.50 nm and a smaller peak at 0.90 nm (Fig. S6D). Palm SS strongly increased the peak height at 0.5 nm, reflecting strong interactions between the Syb2 and Stx1A palmitoyl chains. In contrast, Stx1A^CVCV^ reduced the peak height at 0.50 nm but increased the peak at 0.90 nm. These results suggested that SNARE TMD palmitoylation increases the backbone contact distances between Syb2 C103 and Stx1A C271/C272) by sterically interactions between their respective palmitoyl chains.

The TMD layer (Syb2 T113-Stx1A S281/T282) contact distance distributions showed increased peak positions from 0.74 nm for Stx1A/Syb2 to 1.50 nm, 1.26 nm and 1.94 nm of Palm Stx1A, Palm Syb2 and Palm SS, respectively (Fig. S6C). Similarly, C-terminal residue (Syb2 S115/T116-Stx1A G288) backbone contact distance analysis revealed that the position of the highest peak increased from 1.6 nm of Stx1A/Syb2 to 2.0 nm, 1.8 nm and 2.6 nm of Palm Stx1A, Palm Syb2 and Palm SS, respectively. These shifts are also consistent with the side chain contact distance changes for the TMD layer (Syb2 T113-Stx1A S281/T282) where the peak height at ∼ 0.51 nm was reduced and that at 1.3-1.4 nm was increased by SNARE TMD palmitoylation. Palm SS even shifted the highest peak location from 0.51 nm to 1.84 nm (Fig. S6D). Similar trends are also shown in the C-terminus residue contact analysis. For more details, see Table S14 and S15.

### SNARE TMD palmitoyl chains bend towards the C-termini of the SNARE TMDs in FP simulations

The average time courses of the tilt angles of palmitoyl chains of Syb2, Stx1A, and SNAP-25 showed that they reached a stabilized state during the SNARE complexes zippering after ∼ 500 ns (Fig. S7). The two palmitoyl chains of the Stx1A in Palm Stx1A and Palm SS preferred bending towards the C-terminus of the Stx1A TMD with peak angles of 120° and 110°, with FWHMs of 60° and 50°, respectively. (Fig. S8A). This is comparable to the tilt angles in Stx1A or t-SNARE plasma membrane simulations (Fig. 2D, 4C).

However, the palmitoyl chain of Syb2 also bent towards to the C-terminus of the Syb2 TMD with a peaked of 110° regardless of Stx1A palmitoylation (Fig. S8A). In the Syb2-SV membrane simulations, the Syb2 palmitoyl chain preferred an orientation parallel to the membrane plane, along with similar temporary tilting toward the CP and the IV leaflet, corresponding to the N-terminus and the C-terminus of the Syb2 TMD, respectively (Fig. S1). For more details, see Table S16.

Further RMSF analysis of palmitoyl chain structural fluctuations indicated that the structural fluctuations of all palmitoyl chains were comparable to those observed in the protein or t-SNARE-planar membrane simulations (Fig. S8B). The orientations of the four palmitoyl chains of SNAP-25 were not affected by SNARE TMD palmitoylation. For more details, see Table S17 and S18.

## Discussion

### Stx1A palmitoylation stabilizes upright fusogenic Stx1A conformation

The CGMD simulations presented here revealed that the palmitoyl chains of both, Stx1A and Syb2, localized preferentially at the membrane midplane in planar membrane simulations. In contrast, the palmitoyl chains of SNAP-25 oriented approximately parallel to the membrane normal between the IC leaflets (Fig. 2, 4, S1). Interestingly, Palm Syb2 showed two metastable tilts toward either the CP or the IV leaflets, respectively. While Syb2 palmitoylation did not noticeably affect the tilting dynamics of its TMD, Stx1A palmitoylation significantly decreased the tilt of the Stx1A TMD as well as the tilt of SND and JMD. The changes in tilting were mainly attributed to the Stx1A palmitoylation of the TMD, although the CVCV mutation also induced an albeit smaller change in tilt angle, presumably due to increased hydrophobicity.

The tilt angles of the Palm Stx1A SND, JMD and TMD domains were similar to those of Stx1A in the t-SNARE complex (Figure 3C, 4B). The structure of the t-SNARE complexes was very similar for all variants of Stx1A (Fig. 4). The palm Stx1A assumes a conformation in the membrane that fits the conformation of Stx1A in the tSNARE complex. In contrast, non-palmitoylated Stx1A assumes a conformation where its JMD and SND tilt down, lying flat and adhering to the membrane surface.

These results suggest that two Stx1A palmitoyl chains stabilize the TMD orientation close to the bilayer normal, thereby decreasing the tilt angles of Stx1A’s JMD and SND. This stabilization is expected to facilitate the early stages of priming and spontaneous fusion by facilitating SNARE complex formation (Fig. 7). In fact, the orientation of the Stx1A SND in the t-SNARE complex is ∼ 58° and the angle distribution of Palm Stx1A has also peaked at ∼ 55°. In contrast, the orientation of non-palmitoylated Stx1A rapidly tilts to ∼ 90°, lying down on the membrane surface. Stx1A TMD palmitoylation therefore stabilizes the Stx1A SND conformation that favors SNARE complex formation. Consistent with this result, *in vitro* experiments, have shown that the decreasing Stx1A SND tilt angles favors membrane fusion (25, 26). The stabilization of the Stx1A fusogenic conformation explains the experimental results in ref. (7) showing that inhibiting Stx1A palmitoylation via K260E mutation dramatically reduces spontaneous release. Stx1A^CVCV^ showed also some degree of stabilization of the upright fusogenic conformation, but this stabilization was weaker than that promoted by Stx1A palmitoylation. Consistent with this intermediate behavior, tStx1A^CVCV^ showed a somewhat smaller reduction of spontaneous release experimentally (7).

**Figure 7.**
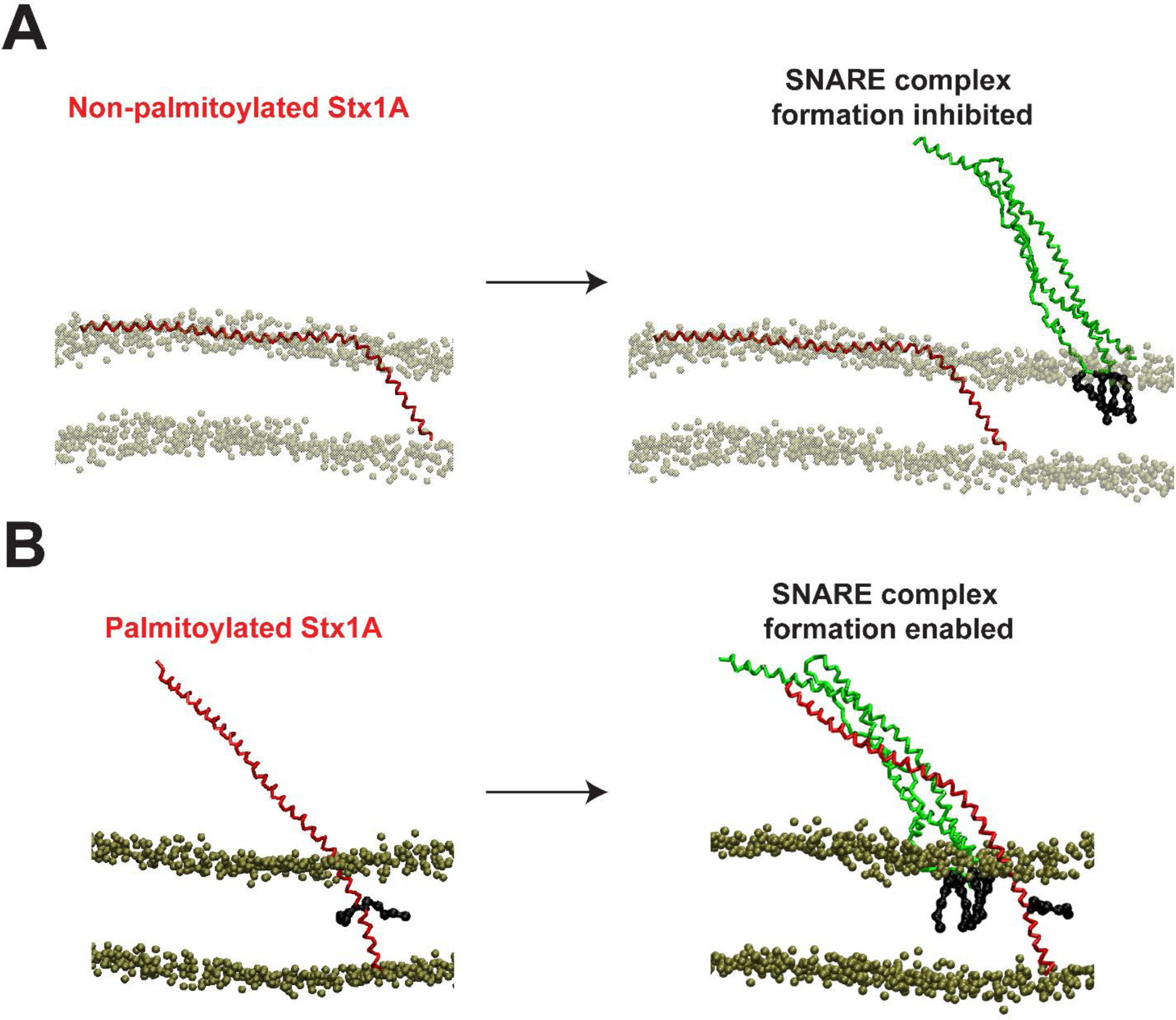
Stx1A C271 and C272 palmitoylation stabilize the Stx1A conformation to facilitate t-SNARE formation. (A) Non-palmitoylation Stx1A (colored red) disfavor binding to SNAP-25 to form a t-SNARE complex by lying flat on the plasma membrane (color brown). (B) Palmitoyl chains of Stx1A stabilize the Stx1A conformation, which facilitates the t-SNARE complex formation and the upstream steps of fusion like docking and priming and spontaneous release.

### Stx1A palmitoylation increases Syb2-Stx1ATMD C-terminal contact distances and delays FP opening

FP simulations showed that SNARE TMD palmitoylation did not affect the final zippering dynamics of the SNARE domain, implying that protein dynamics of SNARE zippering are not crucial in controlling FP formation from the hemifusion diaphragm (HD) state (Fig. 5, S4 and S6A). Notably, simulations with Stx1A TMD palmitoylation exhibited slower distal leaflet lipid transfer and FP opening, suggesting that the dynamics of SNARE TMD interactions beyond zippering of layers 5 to 8 are regulated by SNARE TMD palmitoylation and control the lipid transfer and subsequent FP opening (Fig. 6B, S5, and S6B-D).

From the backbone contact analysis of SNARE TMDs, contacts between cysteine palmitoyl chains of palm Stx1A and palm Syb2 in palm SS, increased their backbone contact distances, as well as backbone contact distances of the TMD layer and between C-terminal residues (Fig. S6B-S6D). This interaction likely stabilizes the HD because the peak location of the distribution of the backbone contact distance between C-terminus residues are 2.1 and 2.6 nm, similar to the minimal head group distance between lipids of IV and EC leaflet of an HD (∼ 2-3 nm), potentially explaining why not all of simulations (6 out of 10) demonstrated distal leaflet lipid transfer and showed FP opening (Fig. 6C). In addition, Stx1A palmitoyl chains also interacted with C103 of Syb2, increasing the contact distances of TMD layer residue contacts as well as C-terminal residue contacts. This effect is weaker for only Syb2 palmitoylation since a closer contact between Palm Syb2 C103 and Stx1A C271 and C272 (the backbone contact distance distribution between cysteine residues also peaked at 0.85 nm) was observed and that contact gets sterically disrupted by two palmitoyl chains of Palm Stx1A (Fig. S6C). Among all SNARE variants, the backbone contact distance between C-terminus residues of Syb2 and Stx1A TMDs is ∼ 3 nm at the mean lag times of these variants, consistent with the observations that FP opened on an HD (Fig. 5 and S4). This suggested that the disruption of C-terminal interactions via TMD palmitoyl chains residue contacts could account for the delay of lag time potentially stabilizing HD.

### Stx1A palmitoylation shortens FP flicker open duration potentially by pushing the distal leaflets to squeeze the FP via palmitoyl chains

FP simulations showed that Stx1A palmitoylation also shortened the flicker open time and elongated flicker closed time (Fig. 6C-F, S5). This may be due to the tilting of the Stx1A palmitoyl chains toward the C-terminus of the Stx1A TMD (the peaked tilt angles are ∼120° and ∼110° for C271 and C272, respectively), which could push the phospholipids at the FP wall near the distal ends, constricting the FP opening. This is also consistent with the reduced open FP conductance changing from 306 ± 21 pS (n = 10) of Stx1A/Syb2 to 243 ± 23 pS (n = 10), even if p = 0.07 from Kruskal-Wallis test. Similar effects were also observed in Syb2 palmitoylation since the tilt angle of C103 is also ∼110°, which also bent toward the C-terminus of Syb2, but the effect is much weaker than Stx1A palmitoylation (from 11 ± 2 ns to 9 ± 2 ns, p = 0.33).

### Bifurcation Effects of Stx1A/Syb2 Dual Palmitoylation on FP Dynamics

Even though single Syb2 palmitoylation has no statistically significant effects altering FP formation (p > 0.05), Palm SS exhibited significant and complex effects on FP dynamics which presumably uncovers a role of Syb2 palmitoylation. Although this variant stabilized the HD and prevented FP formation in 4 out 10 simulations within the 4 µs simulation time by increasing the TMD C-terminus residue contact distance governed by palmitoyl chains, the open FP conductance distribution showed an increase of the peak position from 248 pS to 295 pS, displayed pronounced fluctuations with the FWHM increasing from 258 pS to 312 pS (Fig. 6F). Surprisingly, the typically short FP flicker open time in Stx1A palmitoylation was restored by the additional palmitoylation of Syb2 from 5 ± 1 ns (n = 9) to 14 ± 3 ns (n=5), p = 0.006. This effect may result from interactions between the palmitoyl chains of Syb2 C103 and Stx1A C271 and C272 observed from the contact distance analysis and similar palmitoyl chain tilt angles (Fig. S6D, S8A) Palmitoyl chain of C103 could prevent the palmitoyl chains of Stx1A squeezing the FP, indicated by the statistical test by comparing the RMSF of C271 and C272 with and without Syb2 palmitoylation, potentially preventing FP closure and reversion to the HD state. This may indicate that the function of Syb2 palmitoylation would appear by cooperation of Stx1A palmitoylation. It needs further investigation of the effects of Stx1A/Syb2 dual palmitoylation like how these palmitoylations are regulated at the upstream of the membrane fusion or during membrane fusion. The potential roles for the cooperation between Syb2 and Stx1A palmitoylation are also waiting for uncovering.

### Future directions for FP simulations

Although the FP simulations invested that Stx1A palmitoylation delayed FP opening and shorten FP flicker open time, experimental study in ref. (7) showed that spontaneous fusion is facilitated by Stx1A palmitoylation regulated by Stx1A JMD. This result implies that the regulation (clamping) of spontaneous release presumably occurs at a much earlier upstream stage of the final 4-layer (layer 5-8) zippering and FP formation. Thus, extending the simulations to upstream states and understanding the clamping interaction by synaptotagmin could help understand the regulation of spontaneous release.

## Funding

This work has been supported by US National Institutes of Health (NIH) grants R35GM139608 and R21NS118319.

## Acknowledgements

We thank Frédéric Pincet for critical reading of the manuscript and excellent comments and suggestions.

## Data availability

All files to run simulations, code to extract data from simulation trajectory and code to analyze data and make figures are available in https://zenodo.org/records/14219039, except the simulation trajectory files whose sizes are too large to upload. Their availability is upon request.

## Author contribution

M.L. designed the project, D.A. and S.S. performed the project, D.A. and M.L analyzed data, and all authors wrote and revised the manuscript.

## Declaration of generative AI and AI-assisted technologies in the writing process

During the preparation of this work the first author used ChatGPT in order to improve language and readability. After using this tool/service, the author(s) reviewed and edited the content as needed and took full responsibility for the content of the publication.

## Declaration of interests

The authors declare no competing interests.

## Notes

### Competing Interest Statement

The authors have declared no competing interest.

### Summary of Updates

We updated the movies by shrinking the size of the first 6 movies and uploaded the last 5 movies.

## Reference

1. T. C. Südhof, Neurotransmitter release: the last millisecond in the life of a synaptic vesicle. Neuron 80, 675–690 (2013).

2. J. J. A. R. o. B. Rizo, Molecular mechanisms underlying neurotransmitter release. 51, 377–408 (2022).

3. T. Söllner et al., SNAP receptors implicated in vesicle targeting and fusion. Nature (London) 362, 318–323 (1993).

4. G. R. Prescott, O. A. Gorleku, J. Greaves, L. H. Chamberlain, Palmitoylation of the synaptic vesicle fusion machinery. J Neurochem 110, 1135–1149 (2009).

5. M. Veit, T. H. Sollner, J. E. Rothman, Multiple palmitoylation of synaptotagmin and the t-SNARE SNAP-25. Febs Lett 385, 119–123 (1996).

6. M. Veit, A. Becher, G. J. M. Ahnert-Hilger, C. Neuroscience, Synaptobrevin 2 is palmitoylated in synaptic vesicles prepared from adult, but not from embryonic brain. 15, 408–416 (2000).

7. G. Vardar, A. Salazar-Lazaro, S. Zobel, T. Trimbuch, C. Rosenmund, Syntaxin-1A modulates vesicle fusion in mammalian neurons via juxtamembrane domain dependent palmitoylation of its transmembrane domain. Elife 11 (2022).

8. H. J. Risselada, G. Bubnis, H. J. P. o. t. N. A. o. S. Grubmüller, Expansion of the fusion stalk and its implication for biological membrane fusion. 111, 11043–11048 (2014).

9. H. J. Risselada, H. J. C. o. i. s. b. Grubmüller, How SNARE molecules mediate membrane fusion: recent insights from molecular simulations. 22, 187–196 (2012).

10. H. J. Risselada, C. Kutzner, H. J. C. Grubmüller, Caught in the act: visualization of SNARE-mediated fusion events in molecular detail. 12, 1049–1055 (2011).

11. S. Sharma, M. Lindau, Molecular mechanism of fusion pore formation driven by the neuronal SNARE complex. Proc Natl Acad Sci U S A 115, 12751–12756 (2018).

12. Y. Zhao et al., All SNAP25 molecules in the vesicle-plasma membrane contact zone change conformation during vesicle priming. Proc Natl Acad Sci U S A 121, e2309161121 (2024).

13. J. Rizo, L. Sari, Y. Qi, W. Im, M. M. Lin, All-atom molecular dynamics simulations of Synaptotagmin-SNARE-complexin complexes bridging a vesicle and a flat lipid bilayer. Elife 11 (2022).

14. J. Rizo, L. Sari, K. Jaczynska, C. Rosenmund, M. M. J. P. o. t. N. A. o. S. Lin, Molecular mechanism underlying SNARE-mediated membrane fusion enlightened by all-atom molecular dynamics simulations. 121, e2321447121 (2024).

15. S. J. Marrink, A. H. De Vries, A. E. J. T. J. o. P. C. B. Mark, Coarse grained model for semiquantitative lipid simulations. 108, 750–760 (2004).

16. S. J. Marrink, H. J. Risselada, S. Yefimov, D. P. Tieleman, A. H. de Vries, The MARTINI force field: coarse grained model for biomolecular simulations. The journal of physical chemistry. B 111, 7812–7824 (2007).

17. S. Jo, T. Kim, V. G. Iyer, W. Im, CHARMM-GUI: a web-based graphical user interface for CHARMM. Journal of computational chemistry 29, 1859–1865 (2008).

18. S.-J. Park, N. Kern, T. Brown, J. Lee, W. Im, CHARMM-GUI PDB manipulator: various PDB structural modifications for biomolecular modeling and simulation. J Mol Biol 435, 167995 (2023).

19. L. Monticelli et al., The MARTINI coarse-grained force field: Extension to proteins. Journal of Chemical Theory and Computation 4, 819–834 (2008).

20. M. Mirdita et al., ColabFold: making protein folding accessible to all. Nat Methods 19, 679–682 (2022).

21. S. Sharma, B. N. Kim, P. J. Stansfeld, M. S. Sansom, M. Lindau, A Coarse Grained Model for a Lipid Membrane with Physiological Composition and Leaflet Asymmetry. PloS one 10, e0144814 (2015).

22. A. Lindahl, Hess, & van der Spoel, GROMACS 2021 Manual (Version 2021). Zenodo (2021).

23. A. Stein, G. Weber, M. C. Wahl, R. Jahn, Helical extension of the neuronal SNARE complex into the membrane. Nature 460, 525–528 (2009).

24. W. Humphrey, A. Dalke, K. Schulten, VMD: visual molecular dynamics. Journal of molecular graphics 14, 33–38 (1996).

25. C. Amos et al., Membrane lipids couple synaptotagmin to SNARE-mediated granule fusion in insulin-secreting cells. Mol Biol Cell 35 (2024).

26. V. Kiessling et al., A molecular mechanism for calcium-mediated synaptotagmin-triggered exocytosis. Nat Struct Mol Biol 25, 911-+ (2018).

27. M. L. Waskom, Seaborn: statistical data visualization. Journal of Open Source Software 6, 3021 (2021).

