## Supplemental information for "Syntaxin 1A Transmembrane Domain Palmitoylation Induces a Fusogenic Conformation"

### Supporting methods

#### Simulation performance and visualization

Simulations are performed on Triton cluster from Frost Institute for Data Science and Computing (https://idsc.miami.edu/). All snapshots were taken by VMD (1) and the source files for all movies are available in https://zenodo.org/records/14219039.

#### Simulation trajectory preprocessing

To extract data from simulation trajectories, the coordinates of protein or protein complex for protein-lipid interaction simulations was centered using command *gmx trjconv* and loaded by package *MDTraj* v1.9.3 (2), whereas for FP simulations, nanodisc scaffold protein was centered. Protein and palmitoyl chain tilting dynamics were analyzed by *MDTraj* (2). This is to avoid the discontinuation of the protein representation due to the periodic boundary conditions.

#### Data analysis and visualization

All data analysis figures were made by Python package *matplotlib.pyplot* and *seaborn* (3, 4). All distributions are calculated using kernel-density estimation with parameter cutoff = 0, meaning that the distributions are cut at both ends of the range of the values (https://seaborn.pydata.org/archive/0.11/generated/seaborn.kdeplot.html). The bandwidth of the gaussian kernel was determined using the 'scott' rule (5). The other parameters are the default parameters in *seaborn.kdeplot* function. The peak positions are identified by *scipy.signal.find_peaks* function by comparing the probability of the position with the values of the nearest two points and the full-width at half-maximums (FWHMs) were identified as the positions which are closest but no smaller than the half of the peak value at both sides of the peaks.

#### Supporting Figures and Captions

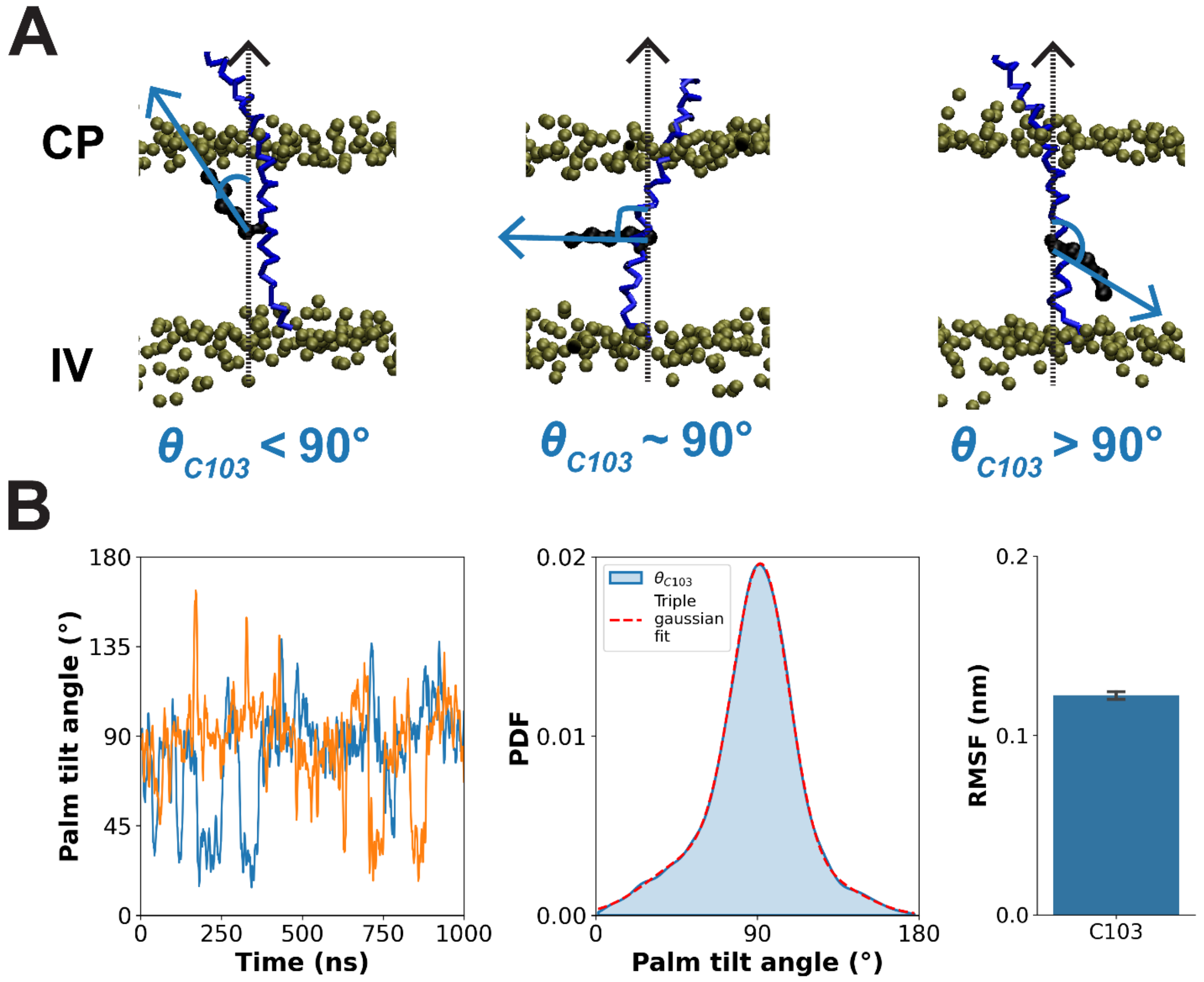


Figure S1. Palmitoyl chain dynamics of Syb2.

(A) Simulation snapshots at 50, 56 and 266 ns that showed the $\theta_{C103} < 90°$, $\theta_{C103} \sim 90°$ and $\theta_{C103} > 90°$, which the palmitoyl chain bent toward the CP leaflet, stayed parallel to the midplane, and bent toward the IV leaflet, respectively. These snapshots were taken from the 1st simulation of the 1st initial condition. The backbones of the Syb2 are colored blue and the head groups of the phospholipids of the SV membrane are colored brown. The tails of the lipids are not shown in these snapshots. (B) The time courses of palmitoyl chain tilting of C103 of Syb2 from 2 simulations (left). They were taken from the 1st simulation of the 1st initial condition (blue) and from the 4th simulation of the 1st initial condition (orange). Due to the huge fluctuations of the raw data, the time courses were preprocessed by applying rolling average with a time window of 5 ns. The distribution of Syb2 palmitoyl chain tilt angle $\theta_{C103}$ does not have a preference to bend toward either the CP or the IV leaflets of the SV membrane (middle). The tilt angle peaked at 91° with the FWHM of 40°. All 10010 frames from all n = 10 simulations were used to calculate the distributions via kernel-density estimation. Because palmitoyl chains occasionally bent toward the CP and the IV leaflet, a triple gaussian function was fit to the distribution with the optimized parameters are $a_1 = 0.003, \mu_1 = 54°, \sigma_1 = 25°, a_2 = 0.018, \mu_2 = 92°, \sigma_2 = 16°, a_3 = 0.002, \mu_3 = 131°$,and $\sigma_3 = 20°$, where $a_i, \mu_i$, and $\sigma_i$ are peak amplitude, peak location and the width of the $i^{th}$ gaussian function (red dash line). The average RMSF of the five side chain beads of palmitoyl chains (right). (0.122 ± 0.002, mean ± SEM).

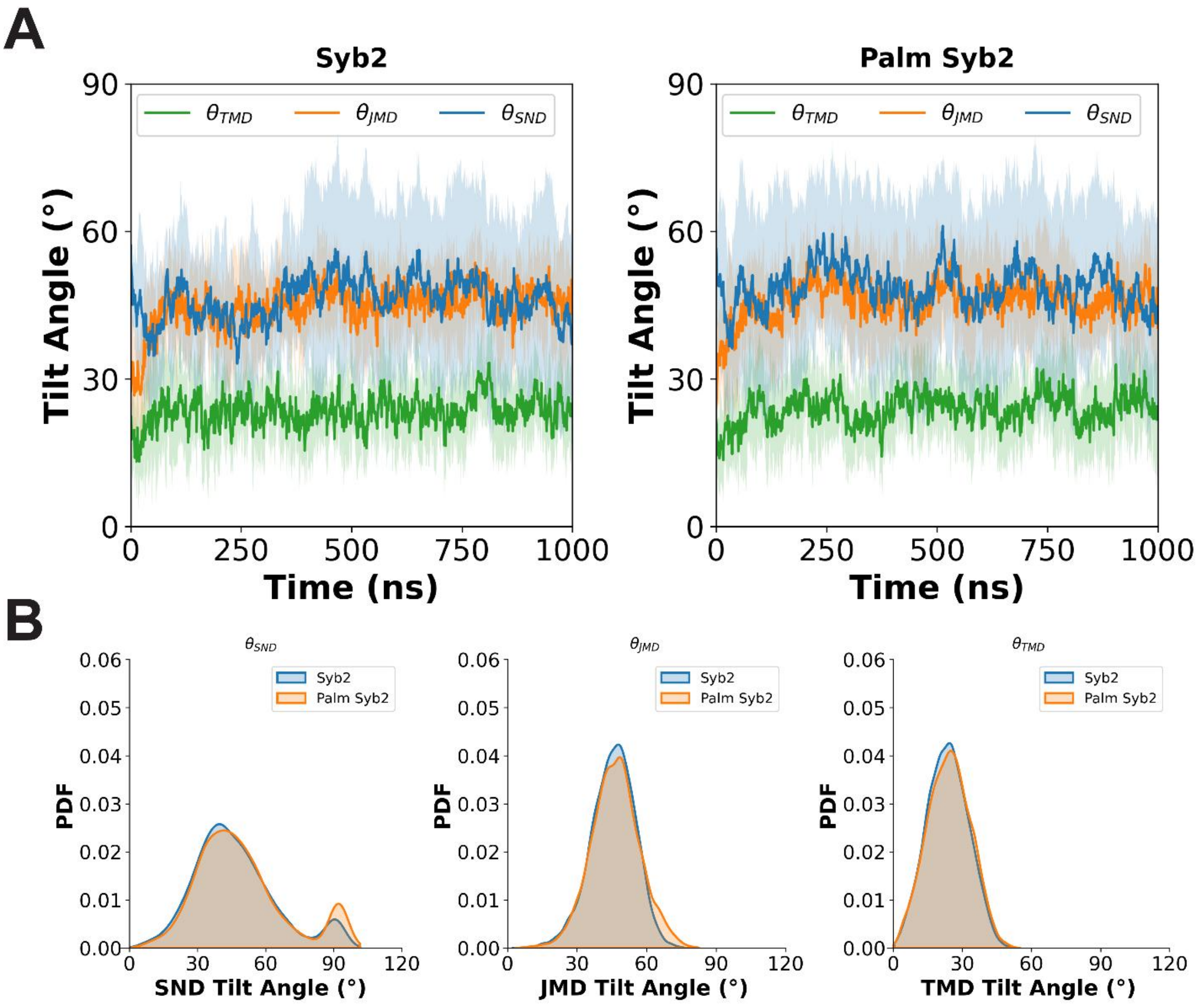


Figure S2. Syb2 palmitoylation does not affect protein tilting dynamics.

(A) The average time courses of $\theta_{\mathrm{SND}}$, $\theta_{\mathrm{JMD}}$ and $\theta_{\mathrm{TMD}}$ for Syb2 and Palm Syb2 are shown (n=10). Shade regions: mean ± SD. (B) The probability density functions (PDFs) of $\theta_{\mathrm{SND}}$, $\theta_{\mathrm{JMD}}$ and $\theta_{\mathrm{TMD}}$ for Syb2 and Palm Syb2 are shown. All frames after 100 ns for all n = 10 simulations were used. Thus, the total numbers of frames are 9010 for non-palmitoylated and palmitoylated Syb2. The Syb2 palmitoylation did not significantly change for tilting dynamics. $\theta_{\mathrm{SND}}$ distribution peaked at 90° and 40° with FWHMs of 12° and 32°, respectively, for Syb2. $\theta_{\mathrm{SND}}$ distribution peaked at 91° and 41° with FWHMs of 12° and 32°, respectively, for Palm Syb2. $\theta_{\mathrm{JMD}}$ distributions for non-palmitoylated and palmitoylated Syb2 both peaked at 48° with an FWHM of 22°. $\theta_{\mathrm{TMD}}$ distributions both peaked at 25° with FWHMs of 22° and 24° for non-palmitoylated Syb2 and palmitoylated Syb2, respectively. For more information, see Table S5.

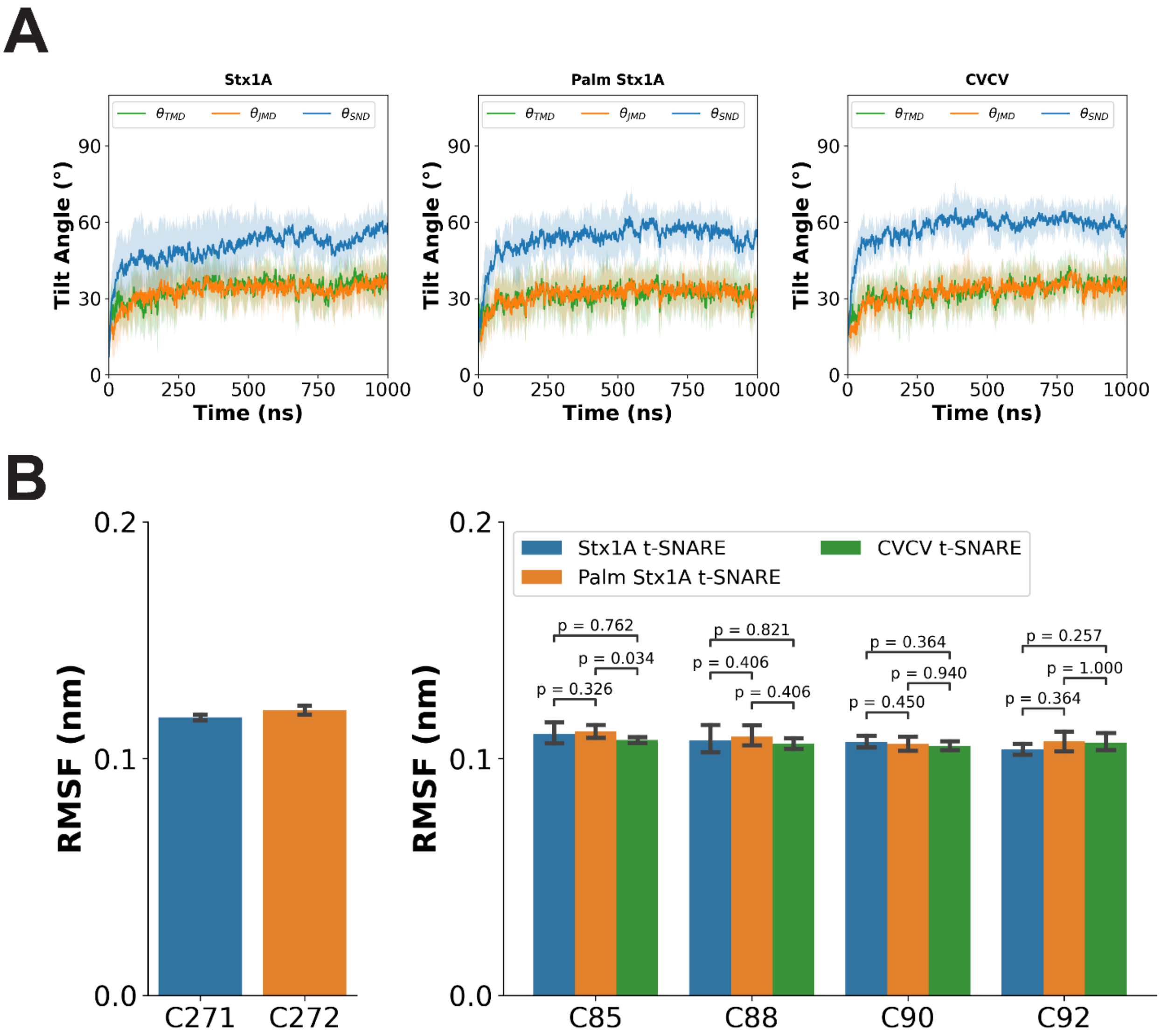


Figure S3. Averaged Stx1A tilting time courses and RMSF of palmitoyl chains.

(A). The average tilt angles time courses of $\theta_{\mathrm{SND}}$, $\theta_{\mathrm{JMD}}$ and $\theta_{\mathrm{TMD}}$ for Stx1A, Palm Stx1A and CVCV Stx1A in t-SNARE complex are shown (n=10). Shade regions: mean ± SD. The tilt angles equilibrated within 500 ns. (B) The RMSFs of palmitoyl chains of C271, C272 of Stx1A, and C85, C88, C90 and C92 of SNAP-25. The structural dynamics of C271 and C272 were similar to single Palm Stx1A in the plasma membrane. The RMSF of C271 and C272 showed a greater structural fluctuation than C85, C88, C90 and C92 of SNAP-25. For more information, see Table S8. Errorbars: SEM and the p-values are from Kruskal-Wallis test.

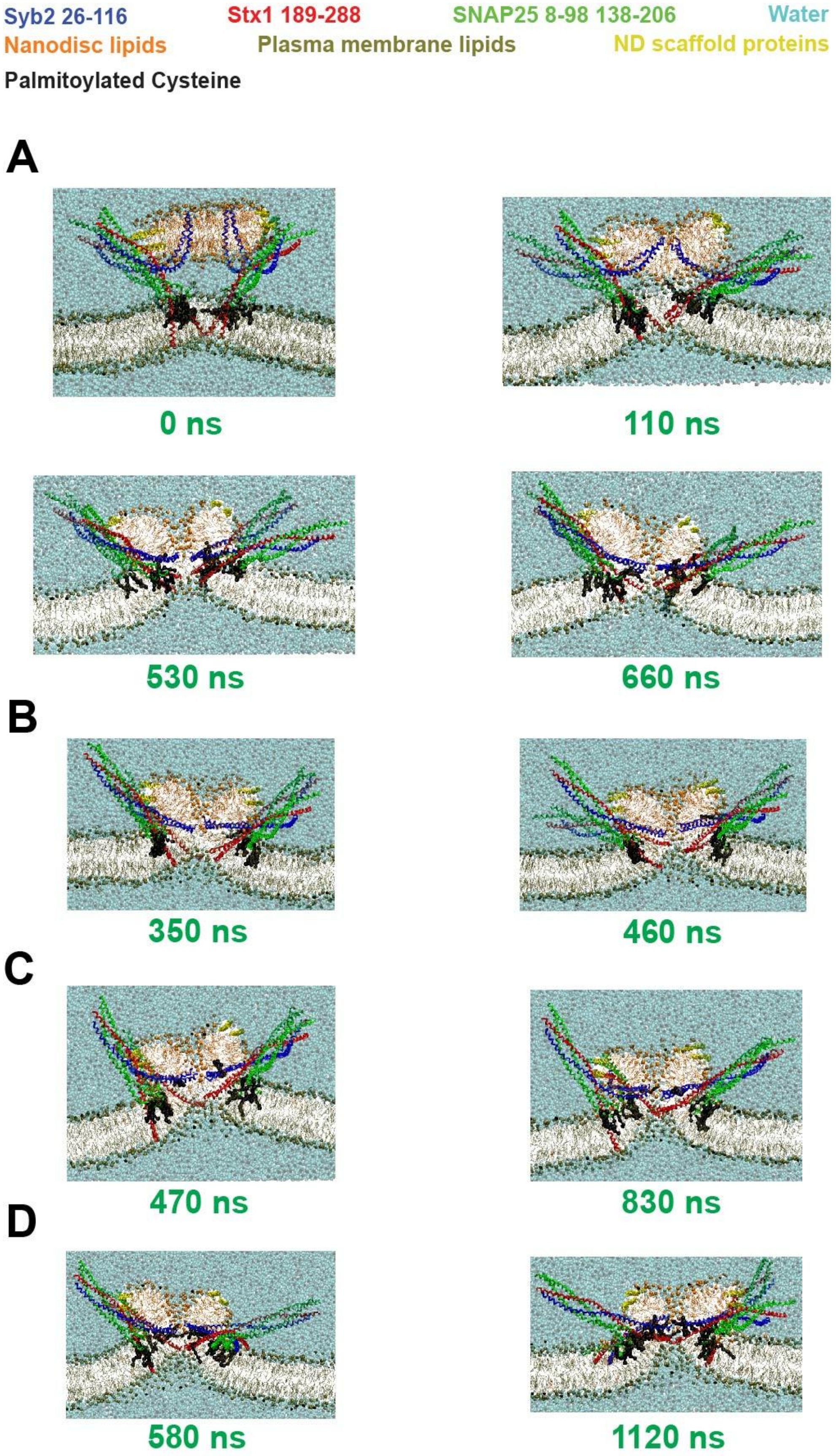
Syb2 26-116
Stx1 189-288
SNAP25 8-98 138-206
Water
Nanodisc lipids
Plasma membrane lipids
ND scaffold proteins
Palmitoylated Cysteine
A
0 ns
110 ns
530 ns
660 ns
B
350 ns
460 ns
C
470 ns
830 ns
D
580 ns
1120 ns

Figure S4. FP open pathways are the same regardless of SNARE TMD palmitoylation.

Snapshots of FP opening for SNARE complexes with Palm Stx1A, CVCV Stx1A, Palm Syb2 and Palm SS. The numbers in green at the bottoms are the simulation time. The backbones of Syb2, Stx1A and SNAP-25 are colored in blue, red and green, respectively. Water beads are colored in cyan transparently. The head groups of nanodisc and plasma membrane phospholipids are colored in orange and brown, respectively. The nanodisc scaffold protein was colored in yellow. (A) FP simulation mediated by Palm Stx1A shared a similar FP open pathway to Fig. 5A-F with not palmitoylated Stx1A. At t = 0, the two membranes stayed away from each other with a distance of ~ 2 nm. At 110 ns, the CP leaflets contacted with the IC leaflet first, and the head groups of the contacted lipids cleared away from the fusion site by forming an ‘X’ like stalk. At 530 ns, HD was formed and the head groups of the phospholipids from the IV leaflet transferred to the EC leaflet at the site that contact Stx1A TMD without FP opening. At 660 ns, the HD disappeared via FP formation and water molecules diffused across the FP. Snapshots were taken from the 3rd simulation. (B) FP open mediated by SNARE complexes with CVCV Stx1A. At 350 ns, HD had been formed and at 460 ns, the FP formed, and water molecules diffused across the FP. Snapshots were taken from the 3rd simulation. (C) FP open mediated by SNARE complexes with Palm Syb2. At 470 ns, HD had been formed and at 830 ns, the FP formed, and water molecules diffused across the FP. Snapshots were taken from the 1st simulation. (D) FP open mediated by SNARE complexes with Palm SS. At 580 ns, HD had been formed and at 1120 ns, the FP formed, and water molecules diffused across the FP. Snapshots were taken from the 5th simulation.

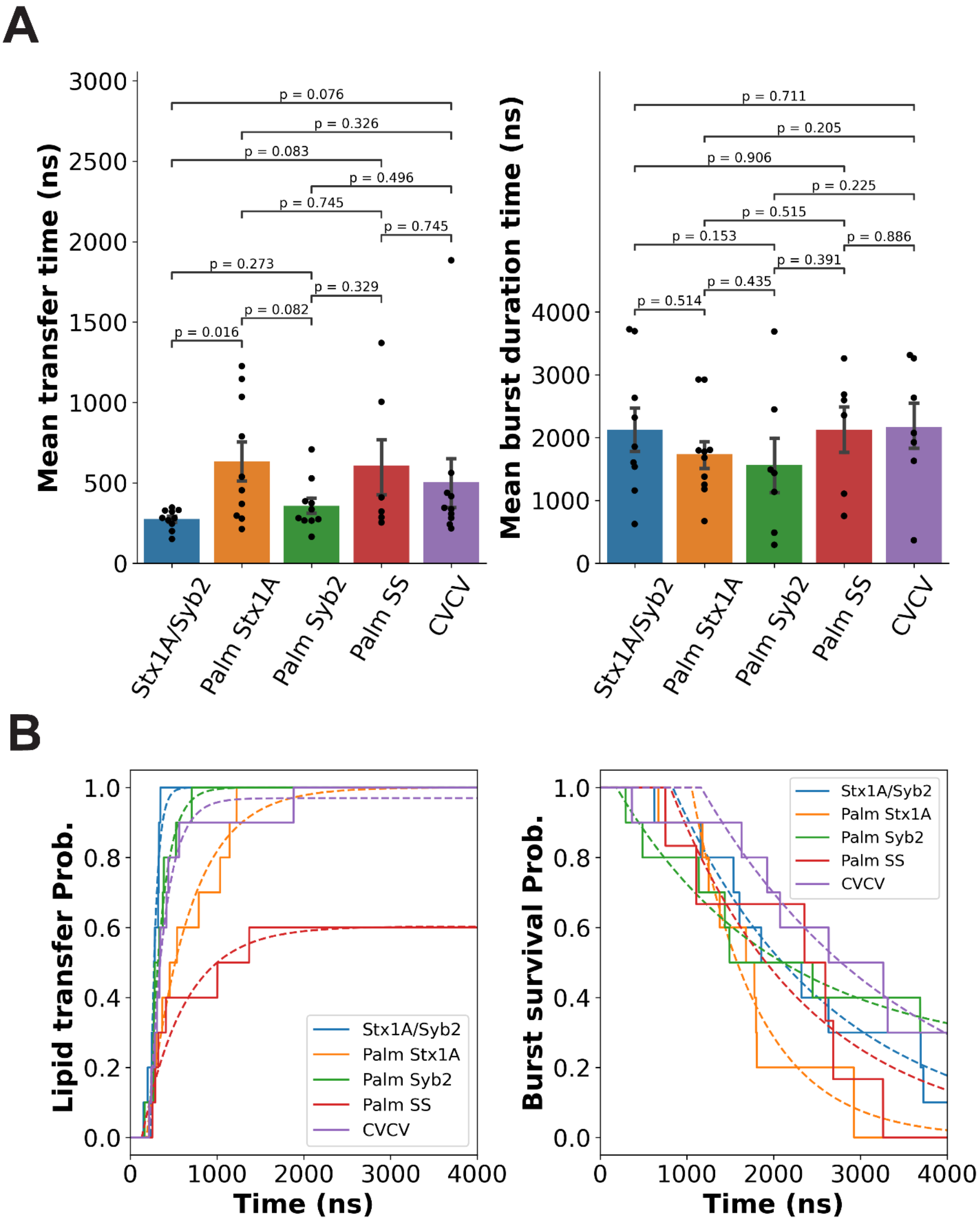


Figure S5. Statistics of distal lipid transfer and burst duration.

(A) The distal leaflet lipid transfer time (left) and burst duration time (right) were measured in FP simulations. Errorbar: SEM. The first 3 digits of p-values for any pair of comparison using Kruskal-Wallis test are shown. The lipid transfer times are Stx1A/Syb2: 276 ± 20 ns, Palm Stx1A: 634 ± 121 ns, Palm Syb2: 358 ± 49 ns, Palm SS: 608 ± 190 ns, and CVCV: 503 ± 157 ns (mean ± SEM, n = 10 except 6 for Palm SS). The burst durations are Stx1A/Syb2: 2128 ± 357 ns, Palm Stx1A: 1739 ± 227 ns, Palm Syb2: 1569 ± 444 ns, Palm SS: 2127 ± 400 ns, and CVCV: 2173 ± 388 ns (mean ± SEM, n = 9, 10, 7, 6 and 7 for these SNARE complexes because the simulations that the burst had not ended within 4 $\mu$s were excluded). The black dots are individual value of each simulation for each variant. (B) The cumulative probability of lipid transfer probability fitted by Eq. 5 and burst survival fitted by Eq. 6 were plotted. Fitted parameters are listed in Table S10, S12. The 4 simulations for Syb2/Stx1A dual palmitoylation that did not show an FP were excluded in these calculations.

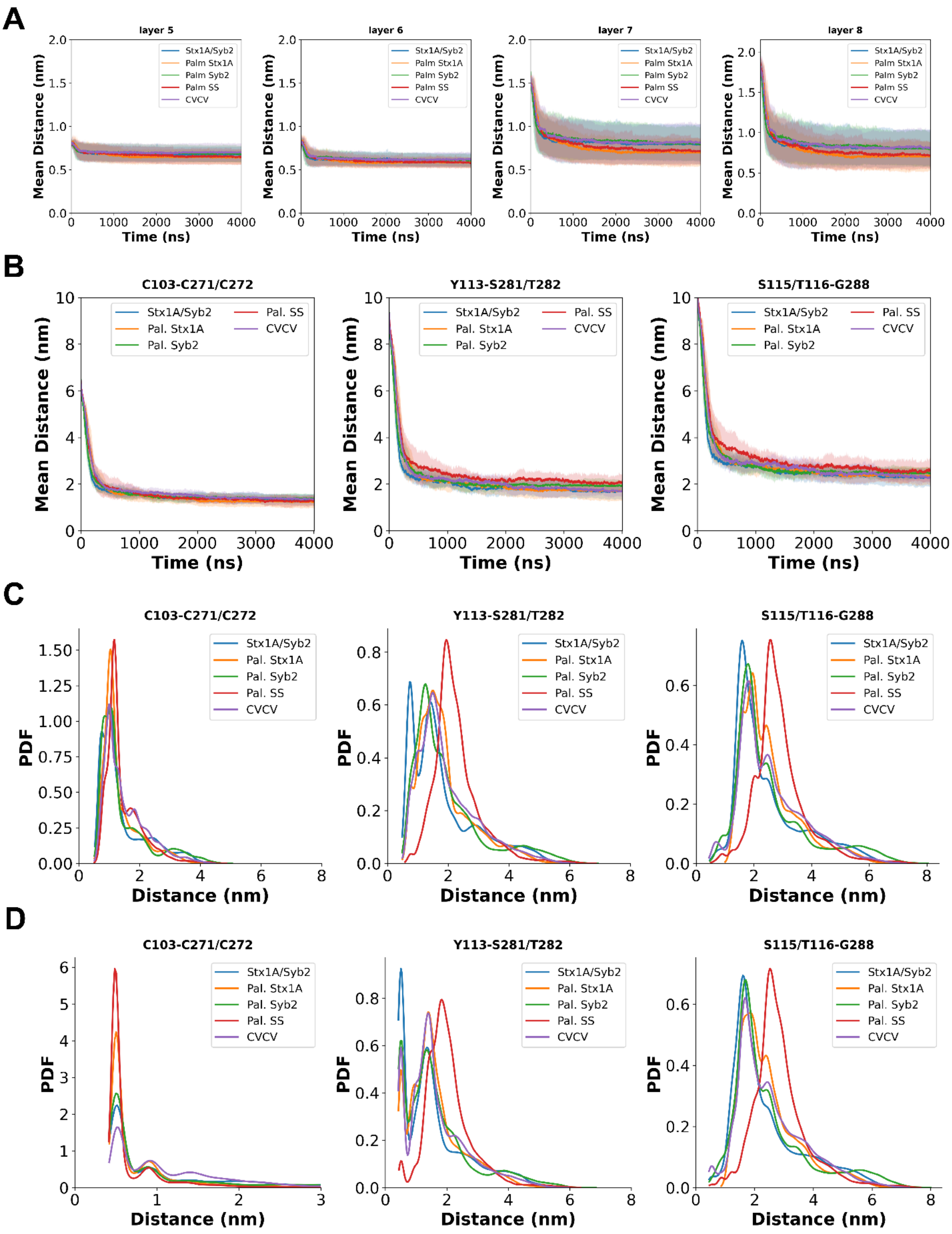


Figure S6. SNARE SND zippering dynamics and TMD contact dynamics in FP simulations.

(A) Average distances between the backbone beads of layer 5-8 residues in Syb2 and Stx1A. All SNARE complexes underwent rapid zippering, with the distance between the backbone beads of these residues stabilizing within < 1 $\mu$s. Shaded region: mean $\pm$ SD. (B) Average distances between the backbone beads of TMD cysteines (C103-C271/C272), hydrophilic layer residues (T113-S281/C272) and C-terminus residues (S115/T116-G288) of Syb2 and Stx1A. The average time courses showed that SNARE TMD palmitoylation and CVCV mutants slowed down the contacts and the Palm SS even results in a larger stable contact distances between the hydrophilic layer and C-terminus residues, which could result in 4 out of 10 simulations failed to nucleate an FP within 4 $\mu$s. At the mean FP lag time in Fig. 6C, the mean contact distance between the backbone beads of C-terminus residues was ~ 3 nm. Shaded region: mean $\pm$ SD. (C) The distribution of the backbone bead contact distance between TMD cysteines (C103-C271/C272) (left), hydrophilic layer residues (T113-S281/C272) (middle) and C-terminus residues (S115/T116-G288) (right) of Syb2 and Stx1A. All 4 SNARE in frames after 500 ns were used. Due to the overlapping of peak locations for SNARE variants, the shades are not plotted. Left: Between TMD cysteine residues, SNARE dual palmitoylation shifted the highest peak location to the larger value from 1.0 nm of Stx1A/Syb2 to 1.2 nm of Palm SS. Palm Stx1A and Palm Syb2 have the highest peaks both at 1.0 nm. The CVCV mutants did not change the location of the highest peak, but generated a new peak at 1.8 nm, like the second highest peak of Palm SS (1.7 nm). Middle: The highest peak location of hydrophilic layer residue backbone contact distance shifted to a higher value from 0.74 nm of Stx1A/Syb2 to 1.5nm, 1.3 nm and 1.94 nm of Palm Stx1A, Palm Syb2 and Palm SS. However, CVCV mutants increased this value to 1.5 nm via weaker C103-V271/V272 interactions than Stx1A/Syb2. Right: Similar trends are also shown in the distribution of C-terminus residue backbone interactions. The highest peak location increased from 1.6 nm of Stx1A/Syb2 to 2.0 nm, 1.8 nm and 2.6 nm for Palm Stx1A, Palm Syb2 and Palm SS. For more details, see Table S13. (D) The distribution of the side chain contact distance between TMD cysteines (C103-C271/C272) (left), hydrophilic layer residues (T113-S281/C272) (middle) and C-terminus residues (S115/T116-G288) (right) of Syb2 and Stx1A. Due to the overlapping of peak locations for SNARE variants, the shades are not plotted. Left: SNARE palmitoylation increased the peak height at 0.50 nm from 2.4 $nm^{-1}$ of Stx1A/Syb2 to 4.2 $nm^{-1}$, 2.6 $nm^{-1}$ and 6.0 $nm^{-1}$ of Palm Stx1A, Palm Syb2 and Palm SS. The CVCV reduced the height of this peak to 1.7 $nm^{-1}$. Middle: The distribution of side chain contact distance of hydrophilic layer residues showed that Palm Stx1A and Palm SS shift the highest peak location from 0.5 nm of Stx1A/Syb2 to 1.4 nm and 1.8 nm, respectively. CVCV also increased the highest peak location to 1.4 nm. But palm Syb2 did not shift the height peak location. Right: The highest peak locations were increased from 1.6 nm of Stx1A/Syb2 to 1.8 nm, 1.7 nm, and 2.5 nm of Palm Stx1A, Palm Syb2 and Palm SS, respectively. CVCV only fine-tuned the highest peak location to 1.7 nm. For more details, see Table S14.

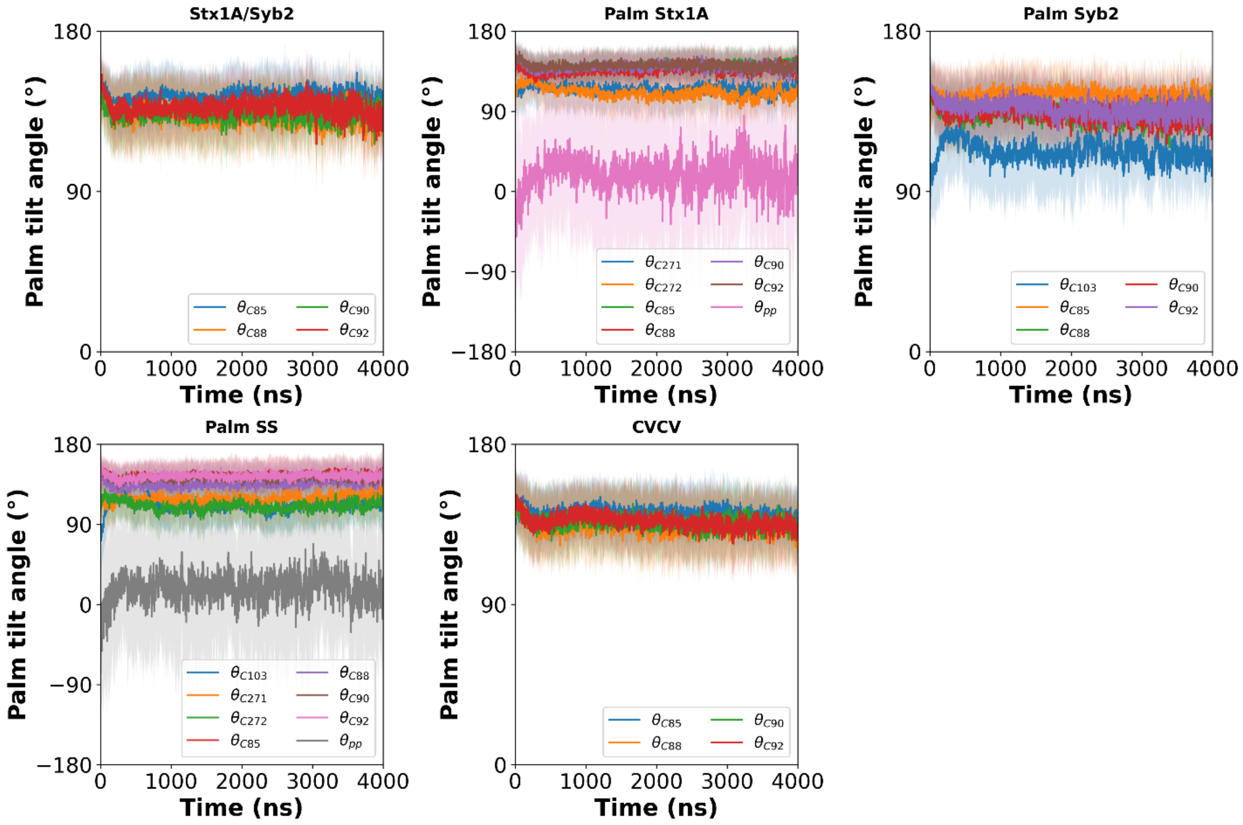


Figure S7. Averaged time courses of palmitoyl chains in SNARE complex.

The tilt angles of palmitoyl chain of C85, C88, C90 and C92 of SNAP-25, C271 and C272 of Stx1A, and C103 of Syb2 were measured. For palmitoylated Stx1A, the angles between the vector of C271 and that of C272 were also measured. All tilt angles are stabilized within 500 ns, which is consistent with the zippering dynamics of SND in SNARE complex.

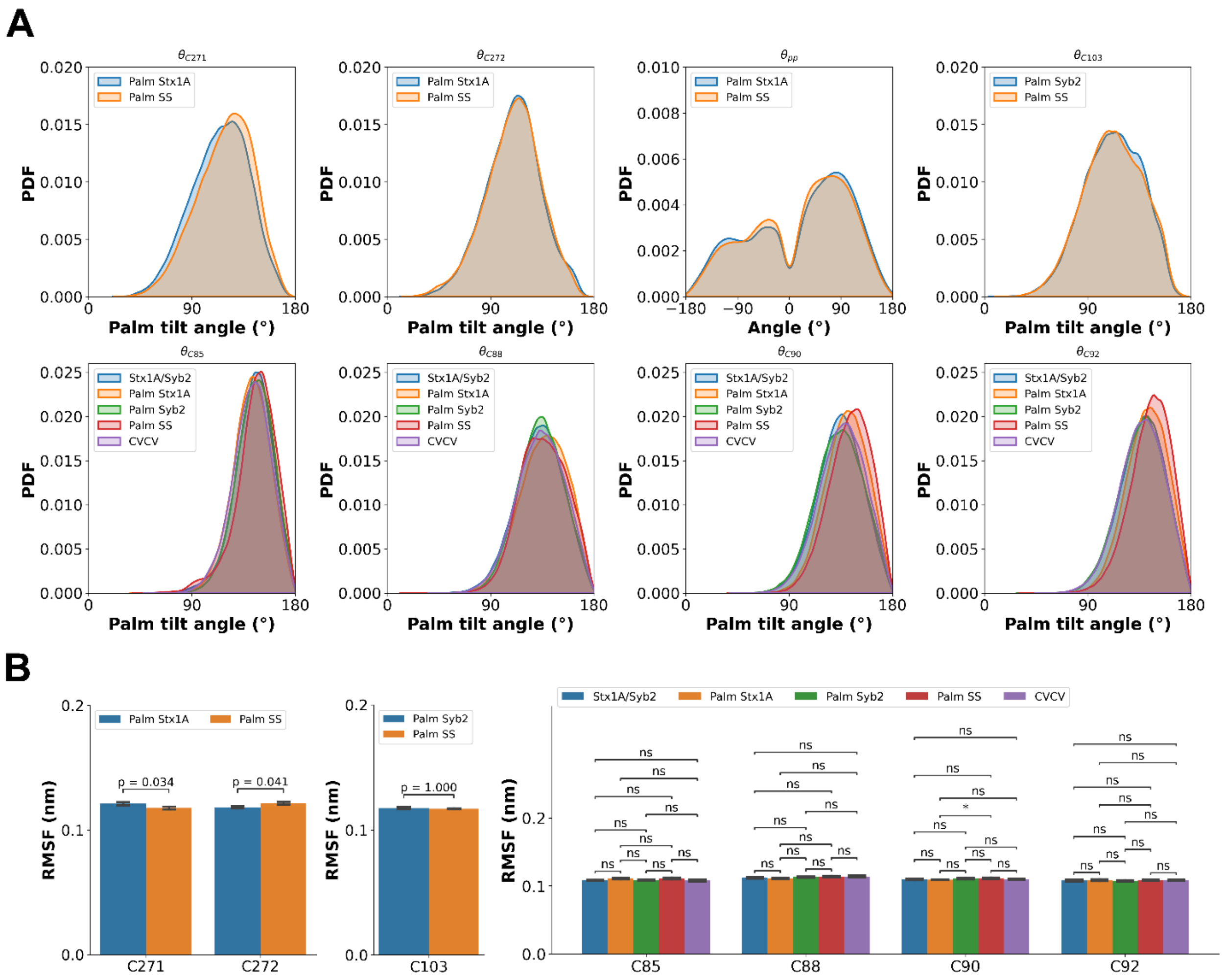


Figure S8. Probability distributions of palmitoyl chain tilting angles and the RMSF of palmitoyl chains in FP simulations.

(A) The distribution of tilt angles of $\theta_{C271}$, $\theta_{C272}$, $\theta_{\mathrm{pp}}$, $\theta_{C103}$, $\theta_{C85}$, $\theta_{C88}$, $\theta_{C90}$, and $\theta_{C92}$ of Palm Stx1A, Palm Syb2 and SNAP-25 in all SNARE systems of FP simulations. All frames after 500 ns were taken to calculate this distribution. The $\theta_{\mathrm{pp}}$ peaked at ~ -100°, ~ -35°, and ~ 85°. The first and the last peaks are similar to the Stx1A and t-SNARE plasma membrane simulations, but FP simulations showed another peak at ~ -35°, which could indicate a transition state during membrane fusion. The Stx1A palmitoylation in Palm Stx1A and Palm SS systems shift the peak angles of $\theta_{C90}$, and $\theta_{C92}$ slightly more parallel to the Stx1A TMD. For more details, please see Table S15. (B) The RMSF of palmitoyl chains of Palm Stx1A, Palm Syb2 and SNAP-25 in all SNARE systems of FP simulations. Dual palmitoylation weakly increased the chain structural fluctuation of C272 of Stx1A but decreased the chain structural fluctuation of C271 of Stx1A. Generally, RMSF of C271, C272 of Stx1A and C103 of Syb2 are ~ 0.12 nm whereas those of C85, C88, C90 and C92 of SNAP-25 are ~ 0.11 nm. For more details about the RMSF and the results of Kruskal-Wallis test, please visit Table S16 and S17, respectively. (*:p $\leq$ 0.05, ns: p > 0.05)

#### Supporting Movies Captions

Movie S1. Palm Stx1A SND relocated to adhere to the plasma membrane and the two palmitoyl chains located to the midplane of the membrane and fluctuated across the membrane but preferred bending to the EC leaflet. In this movie, the Stx1A was centered in the simulation box and water and ions were not shown.

Movie S2. Palm Syb2 SND relocated to adhere to the SV membrane and the palmitoyl chain stay parallel to the midplane of the membrane with fluctuations. In this movie, the Syb2 was centered in the simulation box and water and ions were not shown.

Movie S3. Non-palmitoyl Stx1A SND relocated to adhere to the plasma membrane. In this movie, the Stx1A was centered in the simulation box and water and ions were not shown.

Movie S4. Non-palmitoyl Syb2 SND relocated to adhere to the plasma membrane. In this movie, the Syb2 was centered in the simulation box and water and ions were not shown.

Movie S5. Palm Stx1A palmitoyl chains located to the midplane of the membrane fluctuated across the membrane but preferred bending to the EC leaflet, whereas SNAP-25 palmitoyl chains inserted into the IC leaflet in the t-SNARE complex. In this movie, the t-SNARE complex was centered in the simulation box and water and ions were not shown.

Movie S6. In the not palmitoylated Stx1A complex, SNAP-25 palmitoyl chains inserted into the IC leaflet in the t-SNARE complex. In this movie, the t-SNARE complex was centered in the simulation box and water and ions were not shown.

Movie S7-S11 FP flickering for Stx1A/Syb2 (S7), Palm Stx1A (S8), CVCV (S9), Palm Syb2 (S10) and Palm Stx1A/Syb2 (S11). In movie S7, only lipid head groups and water beads are shown. In movie S8, S10 and S11, in addition to lipid head groups and water beads, the palmitoyl chains of C103, C271 and C272 are shown. In movie S9, in addition to lipid head groups and water beads, the V271 and V272 are shown.

#### Supporting Tables

| Protein or Protein Complex | Membrane size $L^2$ ($nm^2$)[a] | Lipid compositions[b] | Membrane self-assemble time $t_{mb}$ (ns) | System equilibration time (ns) | # self-assembly state | # runs per self-assembly | Simulation time ($\mu$s) |
|---|---|---|---|---|---|---|---|
| Syb2 | $\sim 15 \times 15$ | SV | 200 | 1 | 2 | 5 | 1 |
| Stx1A | $\sim 20 \times 20$ | PM | 500 | 1 | 2 | 5 | 2 |
| Stx1A-SNAP25 | $\sim 20 \times 20$ | PM | 500 | 1 | 2 | 5 | 1 |

Table S1. Hyperparameters of protein-lipid interaction simulations.

Note:

(a). This size refers to the equilibrated membrane size projected on the xy-plane.

(b). SV means that we applied the membrane with the composition in synaptic vesicle listed in ref. (6) whereas PM means that we applied the membrane with the composition in plasma membrane listed in ref. (7).

| Proteins | SND | JMD | TMD |
|---|---|---|---|
| Syb2 | G26-L84 | K85-K94 | M95-T116 |
| Stx1A | K189-V255 | K256-K265 | I266-G288 |

Table S2. The residue ranges of SNARE domains (SND), JMD and TMD of Syb2 and Stx1A in ref. (8).

| Angle[a] | Peak location (°) | Peak value | FWHM (°) | N frames[b] |
|---|---|---|---|---|
| $\theta_{C271}$ | 109 | 0.02 | 50 | 20010 |
| $\theta_{C272}$ | 109 | 0.02 | 40 | 20010 |
| $\theta_{C103}$ | 91 | 0.02 | 40 | 10010 |
| $\theta_{\mathrm{pp}}$ | 84 | 0.005 | 115 | 20010 |
| $\theta_{\mathrm{pp}}$ | -105 | 0.003 | 133 | 20010 |

Table S3. The peak positions and full-width at half-maximums for Palm Stx1A and Palm Syb2 palmitoyl chain tilt angles and angle between the palmitoyl chain of C271 and that of C272 in protein-planar membrane simulations.

Note:

[a]$\theta_{\mathrm{pp}}$ has two peak positions but the peak value for the first one is bigger than the second one.

[b]All frames in all n = 10 simulations were used to calculate the distribution of these angles.

| Stx1A Variants | Angle[a] | Peak location (°) | Peak value | FWHM (°) | N frames[b] |
|---|---|---|---|---|---|
| Not Palm | $\theta_{\mathrm{SND}}$ | 90 | 0.07 | 12 | 2010 |
| Palm | $\theta_{\mathrm{SND}}$ | 91 | 0.014 | 22 | 2010 |
| CVCV | $\theta_{\mathrm{SND}}$ | 86 | 0.02 | 18 | 2010 |
| Palm | $\theta_{\mathrm{SND}}$ | 55 | 0.015 | 46 | 2010 |
| CVCV | $\theta_{\mathrm{SND}}$ | 36 | 0.01 | 83 | 2010 |
| Not Palm | $\theta_{\mathrm{JMD}}$ | 67 | 0.06 | 16 | 2010 |
| Palm | $\theta_{\mathrm{JMD}}$ | 41 | 0.02 | 44 | 2010 |
| CVCV | $\theta_{\mathrm{JMD}}$ | 59 | 0.024 | 42 | 2010 |
| CVCV | $\theta_{\mathrm{JMD}}$ | 39 | 0.019 | 48 | 2010 |
| Not Palm | $\theta_{\mathrm{TMD}}$ | 46 | 0.05 | 18 | 2010 |
| Palm | $\theta_{\mathrm{TMD}}$ | 39 | 0.04 | 21 | 2010 |
| CVCV | $\theta_{\mathrm{TMD}}$ | 42 | 0.05 | 18 | 2010 |

Table S4. The peak positions and FWHMs for Stx1A SND, JMD and TMD tilt angles in protein-planar membrane simulations.

Note:

[a]$\theta_{\mathrm{SND}}$ of Palm Stx1A, CVCV Stx1A and $\theta_{\mathrm{JMD}}$ of CVCV Stx1A have two peaks. The HWHMs of $\theta_{\mathrm{JMD}}$ of CVCV Stx1A are overlapped since the two peaks are very close to each other.

[b]All frames in the last 200 ns of all n = 10 simulations were used to calculate the distribution of these angles.

| Syb2 Variants | Angle[a] | Peak location (°) | Peak value | FWHM (°) | N frames[b] |
|---|---|---|---|---|---|
| Not Palm | $\theta_{\mathrm{SND}}$ | 90 | 0.006 | 12 | 9010 |
| Palm | $\theta_{\mathrm{SND}}$ | 91 | 0.009 | 12 | 9010 |
| Not Palm | $\theta_{\mathrm{SND}}$ | 40 | 0.026 | 32 | 9010 |
| Palm | $\theta_{\mathrm{SND}}$ | 41 | 0.024 | 32 | 9010 |
| Not Palm | $\theta_{\mathrm{JMD}}$ | 48 | 0.04 | 22 | 9010 |
| Palm | $\theta_{\mathrm{JMD}}$ | 48 | 0.04 | 22 | 9010 |
| Not Palm | $\theta_{\mathrm{TMD}}$ | 25 | 0.04 | 22 | 9010 |
| Palm | $\theta_{\mathrm{TMD}}$ | 25 | 0.04 | 24 | 9010 |

Table S5. The peak positions and HWHMs for Syb2 SND, JMD and TMD tilt angles in protein-planar membrane simulations.

Note:

[a]$\theta_{\mathrm{SND}}$ of Syb2 and Palm Syb2 have two peaks.

[b]All frames in the last 900 ns of all n = 10 simulations were used to calculate the distribution of these angles.

| Stx1A Variants | Angle | Peak location (°) | Peak value | FWHM (°) | N frames[a] |
|---|---|---|---|---|---|
| Not Palm | $\theta_{\mathrm{SND}}$ | 57 | 0.05 | 18 | 5010 |
| Palm | $\theta_{\mathrm{SND}}$ | 59 | 0.05 | 16 | 5010 |
| CVCV | $\theta_{\mathrm{SND}}$ | 60 | 0.06 | 14 | 5010 |
| Not Palm | $\theta_{\mathrm{JMD}}$ | 37 | 0.05 | 18 | 5010 |
| Palm | $\theta_{\mathrm{JMD}}$ | 35 | 0.05 | 18 | 5010 |
| CVCV | $\theta_{\mathrm{JMD}}$ | 37 | 0.05 | 18 | 5010 |
| Not Palm | $\theta_{\mathrm{TMD}}$ | 36 | 0.05 | 20 | 5010 |
| Palm | $\theta_{\mathrm{TMD}}$ | 34 | 0.04 | 22 | 5010 |
| CVCV | $\theta_{\mathrm{TMD}}$ | 36 | 0.05 | 20 | 5010 |

Table S6. The peak positions and HWHMs for Stx1A SND, JMD and TMD tilt angles in t-SNARE-planar membrane simulations.

Note:

[a]All frames in the last 500 ns of all n = 10 simulations were used to calculate the distribution of these angles.

| Stx1A Variants | Angle[a] | Peak location (°) | Peak value | FWHM (°) | N frames[b] |
|---|---|---|---|---|---|
| Palm Stx1A | $\theta_{C271}$ | 110 | 0.02 | 45 | 10010 |
| Palm Stx1A | $\theta_{C272}$ | 109 | 0.02 | 42 | 10010 |
| Palm Stx1A | $\theta_{\mathrm{pp}}$ | 87 | 0.005 | 120 | 10010 |
| Palm Stx1A | $\theta_{\mathrm{pp}}$ | -108 | 0.003 | 137 | 10010 |
| Palm Stx1A | $\theta_{C85}$ | 158 | 0.03 | 32 | 10010 |
| Stx1A | $\theta_{C85}$ | 154 | 0.03 | 32 | 10010 |
| CVCV | $\theta_{C85}$ | 160 | 0.03 | 31 | 10010 |
| Palm Stx1A | $\theta_{C88}$ | 161 | 0.03 | 29 | 10010 |
| Stx1A | $\theta_{C88}$ | 161 | 0.03 | 29 | 10010 |
| CVCV | $\theta_{C88}$ | 162 | 0.03 | 28 | 10010 |
| Palm Stx1A | $\theta_{C90}$ | 161 | 0.03 | 28 | 10010 |
| Stx1A | $\theta_{C90}$ | 162 | 0.03 | 28 | 10010 |
| CVCV | $\theta_{C90}$ | 163 | 0.03 | 29 | 10010 |
| Stx1A | $\theta_{C92}$ | 159 | 0.03 | 32 | 10010 |
| Palm Stx1A | $\theta_{C92}$ | 160 | 0.03 | 33 | 10010 |
| CVCV | $\theta_{C92}$ | 160 | 0.03 | 31 | 10010 |

Table S7. The peak positions and FWHM for Stx1A and SNAP-25 palmitoyl chain tilt angles and angle between the palmitoyl chain of C271 and that of C272 in t-SNARE-planar membrane simulations.

Note:

[a]$\theta_{\mathrm{pp}}$ has two peak positions but the peak value for the first one is bigger than the second one.

[b]All frames in all n = 10 simulations were used to calculate the distribution of these angles.

| Stx1A Variants | Palmitoyl Cysteine | Mean RMSF (nm) | SEM (nm) | N simulations |
|---|---|---|---|---|
| Palm Stx1A | C271 | 0.118 | 0.001 | 10 |
| Palm Stx1A | C272 | 0.120 | 0.002 | 10 |
| Stx1A | C85 | 0.110 | 0.002 | 10 |
| Palm Stx1A | C85 | 0.111 | 0.001 | 10 |
| CVCV | C85 | 0.108 | 0.001 | 10 |
| Stx1A | C88 | 0.108 | 0.003 | 10 |
| Palm Stx1A | C88 | 0.110 | 0.002 | 10 |
| CVCV | C88 | 0.106 | 0.001 | 10 |
| Stx1A | C90 | 0.107 | 0.001 | 10 |
| Palm Stx1A | C90 | 0.106 | 0.002 | 10 |
| CVCV | C90 | 0.105 | 0.001 | 10 |
| Stx1A | C92 | 0.104 | 0.001 | 10 |
| Palm Stx1A | C92 | 0.107 | 0.002 | 10 |
| CVCV | C92 | 0.107 | 0.002 | 10 |

Table S8. The RMSF for Stx1A and SNAP-25 palmitoyl chains in t-SNARE-plasma membrane simulations. The structural fluctuations of C271 and C272 palmitoyl chains are similar to that of Palm Stx1A alone and they performed larger structural fluctuations than palmitoyl chains of SNAP-25.

| $\tau_{\text{rise}}$ | $\tau_{\text{delay}}$ | $A$ | MSE | $R^2$ | SNARE Variants |
|---|---|---|---|---|---|
| 85.838 ± 0.297 | 257.849 ± 0.204 | 1.000 ± 0.000 | 0 | 0.998 | Stx1A/Syb2 |
| 1010.607 ± 9.238 | 245.333 ± 3.747 | 0.999 ± 0.002 | 0.003 | 0.971 | Palm Stx1A |
| 186.242 ± 0.934 | 253.573 ± 0.628 | 1.000 ± 0.000 | 0.001 | 0.993 | Palm Syb2 |
| 393.514 ± 3.764 | 290.729 ± 2.327 | 0.601 ± 0.001 | 0.001 | 0.966 | Palm SS |
| 346.931 ± 2.621 | 237.167 ± 1.662 | 0.936 ± 0.001 | 0.002 | 0.977 | CVCV |

Table S9. The fitted parameter of the lag-exponential fitting of FP cumulative burst start probability of FP simulations for WT, palmitoyl Stx1A only, palmitoyl Syb2 only, Stx1-Syb2 double palmitoylation and CVCV mutated Stx1A SNARE complexes.

| $\tau_{\text{rise}}$ | $\tau_{\text{delay}}$ | $A$ | MSE | $R^2$ | SNARE Variants |
|---|---|---|---|---|---|
| 61.037 ± 0.654 | 231.761 ± 0.453 | 1.000 ± 0.000 | 0.001 | 0.985 | Stx1A/Syb2 |
| 477.550 ± 3.100 | 187.083 ± 1.849 | 1.000 ± 0.001 | 0.002 | 0.98 | Palm Stx1A |
| 148.125 ± 0.824 | 215.716 ± 0.560 | 1.000 ± 0.000 | 0.001 | 0.992 | Palm Syb2 |
| 491.317 ± 4.532 | 133.554 ± 2.691 | 0.603 ± 0.001 | 0.001 | 0.953 | Palm SS |
| 179.491 ± 1.784 | 214.198 ± 1.204 | 0.969 ± 0.001 | 0.002 | 0.971 | CVCV |

Table S10. The fitted parameter of the lag-exponential fitting of distal leaflet lipid mixing probability of FP simulations for WT, palmitoyl Stx1A only, palmitoyl Syb2 only, Stx1-Syb2 double palmitoylation and CVCV mutated Stx1A SNARE complexes.

| $\tau_{decay}$ (ns) | $n$ | MSE | $R^2$ | SNARE Variants |
|---|---|---|---|---|
| 2.390 ± 0.011 | 1.231 ± 0.005 | 0 | 0.998 | Stx1A/Syb2 |
| 1.890 ± 0.007 | 1.561 ± 0.008 | 0 | 0.999 | Palm Stx1A |
| 2.337 ± 0.013 | 1.318 ± 0.007 | 0 | 0.997 | Palm Syb2 |
| 2.577 ± 0.021 | 1.092 ± 0.007 | 0 | 0.993 | Palm SS |
| 2.628 ± 0.014 | 1.373 ± 0.008 | 0 | 0.997 | CVCV |

Table S11. The fitted parameter of the power law fitting of flicker open duration of FP simulations for WT, CVCV, palmitoyl Stx1A only, palmitoyl Syb2 only and Stx1-Syb2 double palmitoylation SNARE complexes. The characteristic decay time $\tau_{\mathrm{decay}}$ and the cooperativity $n$ fine-tune the FP flicker open kinetics together.

| $\tau_{decay}$ | $\tau_{delay}$ | $A$ | **MSE** | $R^2$ | **SNARE Variants** |
|---|---|---|---|---|---|
| 1827.118 ± 34.204 | 836.289 ± 5.970 | 1.000 ± 0.009 | 0.003 | 0.97 | Stx1A/Syb2 |
| 759.065 ± 9.667 | 1052.960 ± 4.056 | 1.000 ± 0.003 | 0.004 | 0.972 | Palm Stx1A |
| 1821.450 ± 27.695 | 189.027 ± 6.197 | 0.768 ± 0.005 | 0.002 | 0.958 | Palm Syb2 |
| 1595.697 ± 64.539 | 806.885 ± 13.639 | 1.000 ± 0.017 | 0.017 | 0.866 | Palm SS |
| 2331.310 ± 85.759 | 1162.756 ± 8.935 | 1.000 ± 0.021 | 0.004 | 0.938 | CVCV |

Table S12. The fitted parameter of the lag-exponential fitting of FP burst duration from FP simulations for WT, CVCV, palmitoyl Stx1A only, palmitoyl Syb2 only and Stx1-Syb2 double palmitoylation SNARE complexes.

| SNARE variants | Contact residue pairs | BB distance peak locations (nm) | Peak value ($nm^{-1}$) | N value |
|---|---|---|---|---|
| Syb2/Stx1A | C103-C271/C272 | 0.76 | 0.93 | 87232 |
| Syb2/Stx1A | C103-C271/C272 | 1.02 | 1.10 | 87232 |
| Palm Stx1A | C103-C271/C272 | 1.04 | 1.50 | 103020 |
| Palm Stx1A | C103-C271/C272 | 2.83 | 0.09 | 103020 |
| Palm Syb2 | C103-C271/C272 | 0.85 | 1.04 | 88228 |
| Palm Syb2 | C103-C271/C272 | 1.05 | 1.10 | 88228 |
| Palm SS | C103-C271/C272 | 1.17 | 1.57 | 124224 |
| Palm SS | C103-C271/C272 | 1.70 | 0.39 | 124224 |
| CVCV | C103-V271/V272 | 0.99 | 1.12 | 121972 |
| CVCV | C103-V271/V272 | 1.82 | 0.38 | 121972 |
| Syb2/Stx1A | Y113-S281/T282 | 0.74 | 0.69 | 87232 |
| Syb2/Stx1A | Y113-S281/T282 | 1.43 | 0.61 | 87232 |
| Palm Stx1A | Y113-S281/T282 | 0.78 | 0.29 | 103020 |
| Palm Stx1A | Y113-S281/T282 | 1.50 | 0.65 | 103020 |
| Palm Syb2 | Y113-S281/T282 | 1.26 | 0.68 | 88228 |
| Palm Syb2 | Y113-S281/T282 | 4.47 | 0.07 | 88228 |
| Palm SS | Y113-S281/T282 | 0.77 | 0.04 | 124224 |
| Palm SS | Y113-S281/T282 | 1.94 | 0.85 | 124224 |
| CVCV | Y113-S281/T282 | 1.04 | 0.43 | 121972 |
| CVCV | Y113-S281/T282 | 1.49 | 0.65 | 121972 |
| Syb2/Stx1A | S115/T116-G288 | 1.59 | 0.75 | 87232 |

| | | | | |
|---|---|---|---|---|
| Syb2/Stx1A | S115/T116-G288 | 2.43 | 0.28 | 87232 |
| Palm Stx1A | S115/T116-G288 | 1.61 | 0.51 | 103020 |
| Palm Stx1A | S115/T116-G288 | 1.96 | 0.64 | 103020 |
| Palm Syb2 | S115/T116-G288 | 1.81 | 0.67 | 88228 |
| Palm Syb2 | S115/T116-G288 | 2.41 | 0.34 | 88228 |
| Palm SS | S115/T116-G288 | 2.06 | 0.26 | 124224 |
| Palm SS | S115/T116-G288 | 2.56 | 0.75 | 124224 |
| CVCV | S115/T116-G288 | 1.82 | 0.61 | 121972 |
| CVCV | S115/T116-G288 | 2.48 | 0.36 | 121972 |

Table S13. The peak information of the distribution of backbone bead contact distances of pairs of residues of C103-C271/C272, Y113-S281/T282 and S115/T116-G288 between Syb2 and Stx1A TMD all the four SNARE complexes of FP simulations. All SNARE complex configurations in all n = 10 simulations after 500 ns were used for each variant.

| SNARE variants | Contact residue pairs | Side chain included distance peak locations (nm) | Peak value ($nm^{-1}$) | N value |
|---|---|---|---|---|
| Syb2/Stx1A | C103-C271/C272 | 0.51 | 2.43 | 87232 |
| Syb2/Stx1A | C103-C271/C272 | 0.92 | 0.49 | 87232 |
| Palm Stx1A | C103-C271/C272 | 0.50 | 4.24 | 103020 |
| Palm Stx1A | C103-C271/C272 | 0.89 | 0.72 | 103020 |
| Palm Syb2 | C103-C271/C272 | 0.50 | 2.57 | 88228 |
| Palm Syb2 | C103-C271/C272 | 0.89 | 0.56 | 88228 |
| Palm SS | C103-C271/C272 | 0.49 | 5.96 | 124224 |
| Palm SS | C103-C271/C272 | 0.90 | 0.54 | 124224 |
| CVCV | C103-V271/V272 | 0.51 | 1.65 | 121972 |
| CVCV | C103-V271/V272 | 0.91 | 0.74 | 121972 |
| Syb2/Stx1A | Y113-S281/T282 | 0.51 | 0.93 | 87232 |
| Syb2/Stx1A | Y113-S281/T282 | 1.36 | 0.59 | 87232 |
| Palm Stx1A | Y113-S281/T282 | 0.51 | 0.50 | 103020 |
| Palm Stx1A | Y113-S281/T282 | 1.40 | 0.74 | 103020 |
| Palm Syb2 | Y113-S281/T282 | 0.50 | 0.62 | 88228 |
| Palm Syb2 | Y113-S281/T282 | 1.33 | 0.58 | 88228 |
| Palm SS | Y113-S281/T282 | 0.50 | 0.11 | 124224 |
| Palm SS | Y113-S281/T282 | 1.84 | 0.79 | 124224 |
| CVCV | Y113-S281/T282 | 0.50 | 0.05 | 121972 |
| CVCV | Y113-S281/T282 | 1.39 | 0.74 | 121972 |
| Syb2/Stx1A | S115/T116-G288 | 0.58 | 0.04 | 87232 |

| | | | | |
|---|---|---|---|---|
| Syb2/Stx1A | S115/T116-G288 | 1.60 | 0.69 | 87232 |
| Palm Stx1A | S115/T116-G288 | 1.84 | 0.57 | 103020 |
| Palm Stx1A | S115/T116-G288 | 2.38 | 0.43 | 103020 |
| Palm Syb2 | S115/T116-G288 | 1.70 | 0.68 | 88228 |
| Palm Syb2 | S115/T116-G288 | 2.39 | 0.32 | 88228 |
| Palm SS | S115/T116-G288 | 1.22 | 0.05 | 124224 |
| Palm SS | S115/T116-G288 | 2.53 | 0.72 | 124224 |
| CVCV | S115/T116-G288 | 1.68 | 0.62 | 121972 |
| CVCV | S115/T116-G288 | 2.43 | 0.34 | 121972 |

Table S14. The peak information of the distribution of contact distances including residue side chains of pairs of residues of C103-C271/C272, Y113-S281/T282 and S115/T116-G288 between Syb2 and Stx1A TMD in all the four SNARE complexes of FP simulations. All SNARE complex configurations in all n = 10 simulations after 500 ns were used for each variant.

| Stx1A variants | Angle[a] | Peak location (°) | Peak value | FWHM (°) | N value[b] |
|---|---|---|---|---|---|
| Palm Stx1A | $\theta_{C271}$ | 125 | 0.015 | 63 | 103020 |
| Palm SS | $\theta_{C271}$ | 127 | 0.016 | 58 | 124224 |
| Palm Stx1A | $\theta_{C272}$ | 114 | 0.02 | 50 | 103020 |
| Palm SS | $\theta_{C272}$ | 114 | 0.02 | 51 | 124224 |
| Palm Stx1A | $\theta_{\mathrm{pp}}$ | 82 | 0.005 | 115 | 103020 |
| Palm Stx1A | $\theta_{\mathrm{pp}}$ | -37 | 0.003 | 135 | 103020 |
| Palm Stx1A | $\theta_{\mathrm{pp}}$ | -106 | 0.003 | 146 | 103020 |
| Palm SS | $\theta_{\mathrm{pp}}$ | 75 | 0.005 | 117 | 124224 |
| Palm SS | $\theta_{\mathrm{pp}}$ | -35 | 0.003 | 128 | 124224 |
| Palm SS | $\theta_{\mathrm{pp}}$ | -95 | 0.002 | 301 | 124224 |
| Palm Syb2 | $\theta_{C103}$ | 114 | 0.014 | 68 | 88228 |
| Palm SS | $\theta_{C103}$ | 113 | 0.014 | 69 | 124224 |
| Syb2/Stx1A | $\theta_{C85}$ | 146 | 0.025 | 37 | 87232 |
| CVCV | $\theta_{C85}$ | 145 | 0.024 | 39 | 121972 |
| Palm Stx1A | $\theta_{C85}$ | 142 | 0.025 | 37 | 103020 |
| Palm Syb2 | $\theta_{C85}$ | 148 | 0.025 | 38 | 88228 |
| Palm SS | $\theta_{C85}$ | 150 | 0.025 | 36 | 124224 |
| Syb2/Stx1A | $\theta_{C88}$ | 135 | 0.02 | 50 | 87232 |
| CVCV | $\theta_{C88}$ | 133 | 0.02 | 53 | 121972 |
| Palm Stx1A | $\theta_{C88}$ | 139 | 0.02 | 55 | 103020 |
| Palm Syb2 | $\theta_{C88}$ | 135 | 0.02 | 48 | 88228 |
| Palm SS | $\theta_{C88}$ | 128 | 0.02 | 56 | 124224 |
| Syb2/Stx1A | $\theta_{C90}$ | 136 | 0.02 | 46 | 87232 |
| CVCV | $\theta_{C90}$ | 138 | 0.02 | 49 | 121972 |
| Palm Stx1A | $\theta_{C90}$ | 141 | 0.02 | 47 | 103020 |
| Palm Syb2 | $\theta_{C90}$ | 137 | 0.02 | 52 | 88228 |
| Palm SS | $\theta_{C90}$ | 135 | 0.02 | 56 | 124224 |
| Syb2/Stx1A | $\theta_{C92}$ | 141 | 0.02 | 48 | 87232 |
| CVCV | $\theta_{C92}$ | 140 | 0.02 | 48 | 121972 |
| Palm Stx1A | $\theta_{C92}$ | 144 | 0.02 | 45 | 103020 |
| Palm Syb2 | $\theta_{C92}$ | 141 | 0.02 | 48 | 88228 |
| Palm SS | $\theta_{C92}$ | 148 | 0.02 | 42 | 124224 |

Table S15. The peak positions and FWHM for Palm Syb2, Palm Stx1A and SNAP-25 palmitoyl chain tilt angles and angle between the palmitoyl chain of C271 and that of C272 in FP simulations.

Note:

[a]$\theta_{\mathrm{pp}}$ has two peak positions but the peak value for the first one is bigger than the second one.

[b]All angles in all n = 10 simulations after 500 ns for all 4 SNAREs are joined to calculate the distribution of these angles.

| Stx1A variants | Palmitoyl Cysteine | Mean RMSF (nm) | SEM (nm) | N simulations |
|---|---|---|---|---|
| Palm Stx1A | C271 | 0.121 | 0.001 | 10 |
| Palm SS | C271 | 0.118 | 0.001 | 10 |
| Palm Stx1A | C272 | 0.118 | 0.001 | 10 |
| Palm SS | C272 | 0.121 | 0.001 | 10 |
| Palm Syb2 | C103 | 0.118 | 0.001 | 10 |
| Palm SS | C103 | 0.117 | 0.0005 | 10 |
| Syb2/Stx1A | C85 | 0.108 | 0.001 | 10 |
| CVCV | C85 | 0.108 | 0.001 | 10 |
| Palm Stx1A | C85 | 0.111 | 0.001 | 10 |
| Palm Syb2 | C85 | 0.109 | 0.001 | 10 |
| Palm SS | C85 | 0.111 | 0.001 | 10 |
| Syb2/Stx1A | C88 | 0.112 | 0.001 | 10 |
| CVCV | C88 | 0.114 | 0.001 | 10 |
| Palm Stx1A | C88 | 0.111 | 0.001 | 10 |
| Palm Syb2 | C88 | 0.113 | 0.001 | 10 |
| Palm SS | C88 | 0.114 | 0.0005 | 10 |
| Syb2/Stx1A | C90 | 0.109 | 0.001 | 10 |
| CVCV | C90 | 0.110 | 0.0004 | 10 |
| Palm Stx1A | C90 | 0.109 | 0.0004 | 10 |
| Palm Syb2 | C90 | 0.111 | 0.001 | 10 |
| Palm SS | C90 | 0.112 | 0.001 | 10 |
| Syb2/Stx1A | C92 | 0.108 | 0.001 | 10 |
| CVCV | C92 | 0.108 | 0.001 | 10 |
| Palm Stx1A | C92 | 0.109 | 0.001 | 10 |
| Palm Syb2 | C92 | 0.108 | 0.001 | 10 |
| Palm SS | C92 | 0.109 | 0.001 | 10 |

Table S16. The RMSF for Stx1A, Syb2 and SNAP-25 palmitoyl chains in FP simulations.

| SNARE variant pairs | C85 | C88 | C90 | C92 |
|---|---|---|---|---|
| Stx1A/Syb2-CVCV | 0.7055 | 0.1988 | 0.1736 | 0.4963 |
| Stx1A/Syb2-Palm Stx1A | 0.1124 | 0.6501 | 0.5967 | 0.7055 |
| Stx1A/Syb2-Palm Syb2 | 0.6501 | 0.2899 | 0.2899 | 0.8206 |
| Stx1A/Syb2-Palm SS | 0.1306 | 0.1988 | 0.0963 | 0.5967 |
| CVCV-Palm Stx1A | 0.1124 | 0.1124 | 0.1736 | 1 |
| CVCV-Palm Syb2 | 0.4963 | 0.7055 | 0.8798 | 0.6501 |
| CVCV-Palm SS | 0.1124 | 0.8798 | 0.2899 | 0.8798 |
| Palm Stx1A-Palm Syb2 | 0.2899 | 0.1306 | 0.2899 | 0.7055 |
| Palm Stx1A-Palm SS | 0.8798 | 0.1509 | 0.0494 | 0.9397 |
| Palm Syb2-Palm SS | 0.2568 | 0.9397 | 0.4497 | 0.5967 |

Table S17. P-values for pair of comparison of SNAP-25 palmitoyl chain RMSF between pairs of SNARE complex states.

### Reference

1. W. Humphrey, A. Dalke, K. Schulten, VMD: visual molecular dynamics. *Journal of molecular graphics* **14**, 33-38 (1996).
2. R. T. McGibbon *et al.*, MDTraj: a modern open library for the analysis of molecular dynamics trajectories. *Biophysical journal* **109**, 1528-1532 (2015).
3. M. L. Waskom, Seaborn: statistical data visualization. *Journal of Open Source Software* **6**, 3021 (2021).
4. J. D. Hunter, Matplotlib: A 2D graphics environment. *Computing in science & engineering* **9**, 90-95 (2007).
5. D. W. Scott, *Multivariate density estimation: theory, practice, and visualization* (John Wiley & Sons, 2015).
6. S. Sharma, B. N. Kim, P. J. Stansfeld, M. S. Sansom, M. Lindau, A Coarse Grained Model for a Lipid Membrane with Physiological Composition and Leaflet Asymmetry. *PloS one* **10**, e0144814 (2015).
7. S. Sharma, M. Lindau, Molecular mechanism of fusion pore formation driven by the neuronal SNARE complex. *Proc Natl Acad Sci U S A* **115**, 12751-12756 (2018).
8. A. Stein, G. Weber, M. C. Wahl, R. Jahn, Helical extension of the neuronal SNARE complex into the membrane. *Nature* **460**, 525-528 (2009).
